# CRISPR-Cas9 Targeting of G-Quadruplex DNA in ADH1 promoter Highlights its role in Transcriptome and Metabolome Regulation

**DOI:** 10.1101/2025.03.14.643214

**Authors:** Ikenna Obi, Pallabi Sengupta, Nasim Sabouri

## Abstract

G-quadruplex (G4) structures are critical regulators of gene expression, yet the role of an individual G4 within its native chromatin remains underexplored. Here, we used CRISPR-Cas9 to introduce guanine-to-thymine mutations at a G4-forming motif within the *adh1^+^* promoter in yeast, creating two mutant strains: one with G4-only mutations and another with both G4 and TATA-box mutations. Chromatin immunoprecipitation using BG4 antibody confirmed reduced G4 enrichment in both mutants, validating G4 structure formation in the wild-type chromatin. Detailed characterizations demonstrated that the G4 mutations alter its dynamics without fully preventing its formation. These mutations significantly reduce *adh1* transcript levels, with G4 TATA-box mutant causing the strongest transcriptional suppression. This indicates a positive regulatory role for the Adh1 G4 structure in *adh1^+^* gene expression. Furthermore, both mutants displayed altered transcriptomic profiles, particularly impacting the oxidoreductase pathway. Metabolomic analyses by mass spectrometry further highlighted substantial disruptions in NAD+/NADH metabolism, a key energy reservoir for metabolic regulation. Together, our findings illustrate how deregulation of a single G4 structure influences transcriptome regulation, with implications for metabolic diseases. It also highlights the therapeutic potential of G4 modulation as a novel, controlled approach to reprogram cellular metabolism to achieve targeted phenotypic shifts.

**GRAPHICAL ABSTRACT:** 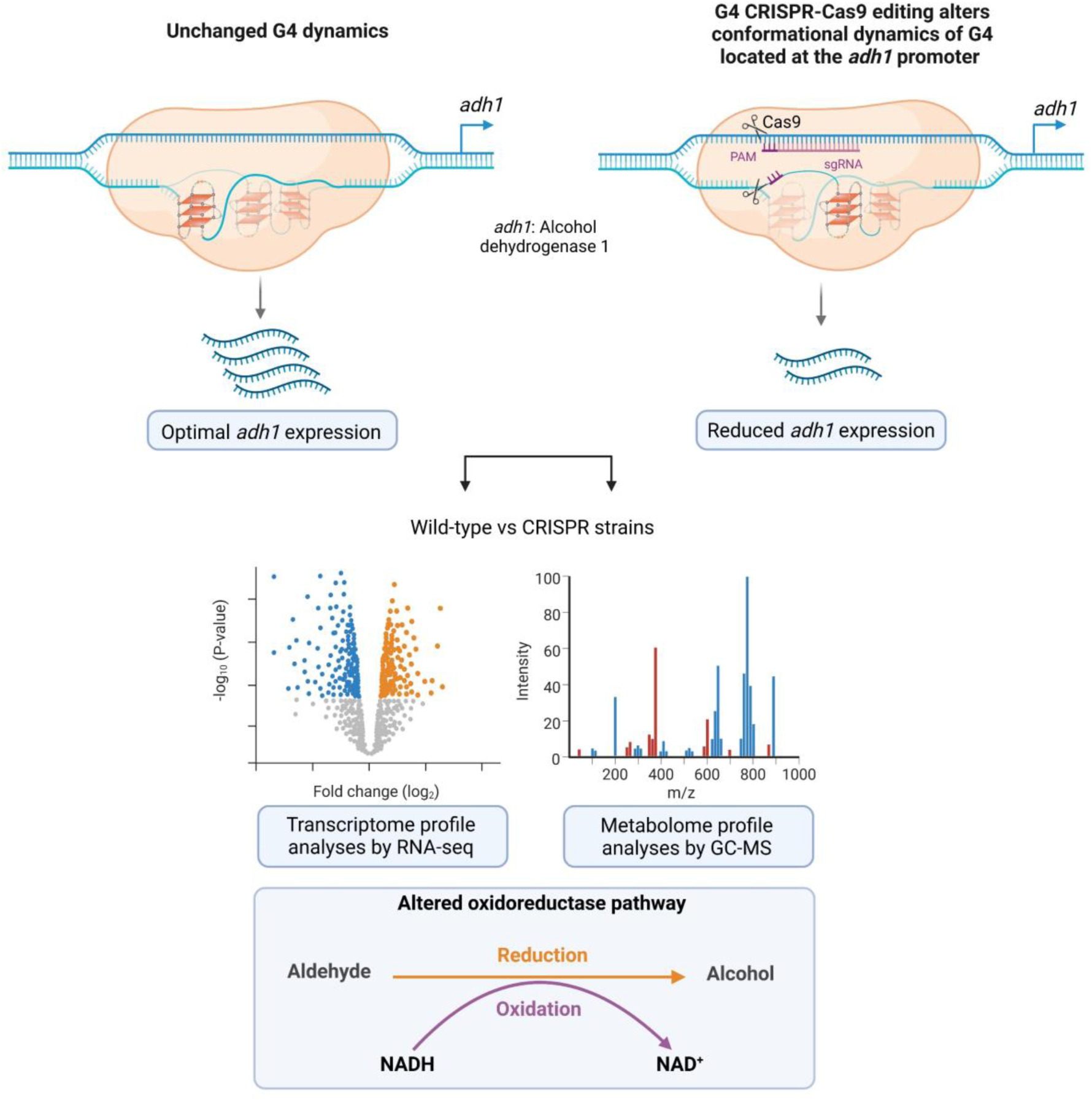

## INTRODUCTION

G-quadruplexes (G4s) are stable non-canonical secondary DNA and RNA structures, formed by tandem guanine (G) repeats when nucleic acids are in their single-stranded state (1). Unlike classical Watson-Crick DNA base pairing observed in B-DNA (2), G4 DNA involves Hoogsteen hydrogen bonding of four G bases to form a G-quartet (3). Two or more G-quartets that stack upon each other form a G4 structure, which is further stabilized by central potassium or sodium ions (4). Techniques or/and tools such as bioinformatic analyses (5–11), G4-sequencing (12,13), selective G4 ligands (14–23), and G4-specific antibodies (24–28) have demonstrated that G4s are prevalent in genomic regions like nucleosome-depleted or open chromatin regions, as well as in replication origins, gene promoters, cis-regulatory elements, untranslated regions (UTRs), and telomeres (9,29–43). These findings suggest that G4 structures impact diverse biological processes (5,8,9,39,44–52). While such systematic studies have significantly advanced our understanding of G4s as a collective phenomenon, they fail to capture the unique contributions of individual G4s within their native chromatin contexts.

Evolutionary conservation of G4 motifs at promoters implies their importance in gene regulation (45), yet dissecting the functional significance of individual G4s is critical for a mechanistic understanding of their roles. The CRISPR-Cas9 system offers a programmable tool for precise gene editing without introducing external tags (53,54), making it ideal for investigating individual G4s. The system requires a specific sequence, known as the protospacer adjacent motif (PAM), to recognize and bind to target DNA. In G4-rich regions, the presence of multiple NGGN sequences (N, any nucleotide) results in numerous PAM sites spaced closely together. This abundance complicates precise targeting by CRISPR-Cas9, leading to off-target effects and reduced specificity. Recent innovations, such as CRISPR approaches using modified Cas9 proteins (55,56) or fusions with G4-binding proteins like nucleolin, have improved specificity (56). Notably, the Balasubramanian lab demonstrated the feasibility of CRISPR-based modification of the endogenous MYC G4 structure, highlighting the promise of such strategies for uncovering the distinct functions of G4s in their native chromatin environments (57). This innovative approach demonstrates the potential use of CRISPR to elucidate the functions of individual G4s within their native chromatin contexts. However, significant challenges remain in comprehensively understanding the functions of all G4s particularly due to limitations in G4 detection technologies (58) and the complexity of higher eukaryotes, such as the human genome, which contains approximately 700,000 putative G4 structures (13).

Yeast genetics has brought powerful approaches for investigating biological questions in eukaryotic organisms, and recently, genetic mutations in fission yeast, *Schizosaccharomyces pombe,* using the CRISPR-Cas9 system were described (59–63). One of the key advantages of using *S. pombe* as a model organism for G4 biology is its significantly smaller genome and thereby the smaller number of G4 motifs—approximately 450 (9) — compared to the ∼700,000 (12) putative G4 structures in the human genome. *S. pombe,* often referred to as the “micro-mammal”, shares many features of chromosome dynamics with higher eukaryotes (64,65). In fact, like humans, *S. pombe* carries G4 motifs at evolutionary conserved genomic features, such as at promoters, telomeres, nucleosome-depleted regions, UTRs, and ribosomal DNA (9,13). Stabilization of G4 structures by G4-selective small molecules causes growth defects during S-phase and slows replication fork progression, indicating that stable G4s present barriers to DNA replication (66,67). In our previous study, we employed chromatin immunoprecipitation combined with sequencing (ChIP-seq) to map G4 structures in *S. pombe* (67,68). We identified approximately 100 stable putative G4 structures out of the 450 G4 motifs, which impeded replication fork progression (67,68). Further, *in vitro* experiments confirmed the formation and topologies of stable G4 structures within several of these identified G4 motifs (67,69–71). The biological significance of these structures is further emphasized by the roles of members of the Pif1 and DEAD-box helicases (72–74) that directly interact with and regulate G4 structures (69,74–77), as well as promoting replication through G4 motifs (9,68,73). These findings demonstrate the conserved roles of G4s and underscore the value of *S. pombe* as a powerful model organism for dissecting the functional roles of individual G4 structures, addressing current limitations in G4 biology.

Building on these findings, we designed a G4 CRISPR-Cas9 mutational approach in *S. pombe* to investigate the functional role of a specific G4 motif. This motif is located upstream of the *adh1^+^*gene, which encodes alcohol dehydrogenase 1 (Adh1), an abundant zinc-binding oxidoreductase required for the reduction of acetaldehyde to ethanol. By targeting this G4 motif, we aimed to address current gaps in understanding the regulatory roles of individual G4 structures in gene expression and metabolic processes. Using the CRISPR-Cas9 system, we simultaneously substituted G residues with thymine (T) residues in four G-tracts of the G4 motif. Bioinformatics analyses and detailed *in vitro* characterizations of the mutated Adh1 G4 sequences confirmed their reduced propensity to form G4 structures. To investigate G4 structure formation in native chromatin, we performed chromatin immunoprecipitation (ChIP) in *S. pombe* cells using the G4-specific BG4 antibody, which identifies a wide range of G4 topologies. We found a significant enrichment of G4 structures at the G-rich motif upstream of the *adh1^+^* gene. Additionally, qPCR stop assays on genomic DNA demonstrated that the wild-type Adh1 G4 (ADH1 G4 WT) motif effectively stalled DNA synthesis, indicating its ability to form stable G4 structures. Altering the ADH1 G4 WT structure reduced *adh1^+^* gene expression, highlighting the importance of the WT G4 structure for optimal gene regulation. Furthermore, disrupting the WT G4 structure led to dysregulation of global cellular transcription and metabolic profiles, underscoring the broader impact of a single G4 structure. These findings illustrate how mutations in an individual G4 motif can disrupt cellular metabolic processes, potentially elevating the risk of metabolic diseases.

## MATERIALS AND METHODS

### Strains

The bacterial and yeast strains used in this study are listed in Supplementary Table 1.

### Oligonucleotides

The oligonucleotides used in this study are shown in Supplementary Table 2.

### Cloning of sgRNA into pLSB-NAT plasmid

pLSB-NAT plasmid with the *S. pombe* codon-optimized Cas9 gene (62) was a generous gift from Robin Allshirés laboratory (University of Edinburgh, (Addgene plasmid # 166698; http://n2t.net/addgene:166698 ; RRID:Addgene_166698). The optimized protocol for cloning sgRNA into pLSB-NAT plasmid is described in (62). Briefly, two different sgRNA were designed, sgRNA1 and sgRNA2. The primers for sgRNA1 or sgRNA2 (Supplementary Figure 2A) targeting G4 motif on antisense strand upstream of *adh1^+^* gene were designed using CRISPR4P primer design tool (http://bahlerweb.cs.ucl.ac.uk/cgi-bin/crispr4p/webapp.py) (60).

By flanking the designed sgRNA sequences with 5′ Ctaga*GGTCTC*gGACT and 3′ GTTTcGAGACCcttCC sequences that contain BsaI site, a 52-mer oligonucleotide DNA sequence was formed, termed as sg1 G435 fw, sg1 G435 rev, and sg2 G435 fw, sg2 G435 rev, respectively (Supplementary Table 2). The sense and antisense DNA of the 52-mer oligonucleotide were hybridized by heating at 95 °C and slowly cooling down to room temperature. Using a Golden Gate reaction (NEB Golden Gate assembly kit), the newly hybridized duplex oligonucleotide was ligated into pLSB-NAT plasmid. *E. coli* cells were transformed with the ligated plasmid and selected on LB + ampicillin agar plates. Colonies without green fluorescence were selected for miniprep plasmid isolation given that BsaI treatment removes the GFP fragment upon successful integration of sg1 G435 or sg2 G435 inserts. In addition, the plasmids from these colonies were digested with NcoI to identify positive clones, which show two fragment sizes of about 4.3 kb and 6.3kb while negative clones will show two fragment sizes of about 5.3 kb and 6.3kb. Finally, the integration of the two different sgRNAs, sgRNA1, and sgRNA2, in pLSB-NAT plasmids was confirmed by sequencing.

### Generation of HR templates

The HR templates were designed such that they have at least 77 base pairs (bp) target homology on each side from the PAM sequence. In addition, the HR templates contain mutations that disrupt the PAM sequence to prevent repeated DNA cleavages. Three pairs of oligonucleotides constituting sense and antisense sequences were used to generate each HR template (HR1 and HR2). Each oligonucleotide pair has 20 bp overlap with the adjacent oligonucleotide pair. The sizes of the oligonucleotide pairs were 74, 97 and 102 bp (Supplementary Table 2). After hybridization of each oligonucleotide pair as described above, recombinational PCR was performed to fuse all three pairs together forming a 233 bp DNA fragment.

### Generation of *S. pombe* strains carrying mutations in the G4 motif

By electroporation, YNS119, a haploid *S. pombe* strain was transformed with sgRNA1 or sgRNA2 integrated into pLSB-NAT and HR1 or HR2, respectively. The cells were plated on EMM2 media (Formedium) containing nourseothricin (Werner BioAgents GmbH) to select for the successful transformants containing the pLSB-NAT plasmids and grown at 30 °C for five days. Transformants were re-streaked on non-selective EMM2 media to facilitate the loss of the pLSB-NAT plasmid. Colony PCR was performed to amplify a region containing the upstream Adh1 G4 motif resulting in 233 bp DNA fragment. The mutated ADH1 G4 motif contains Eae1 restriction site (G*TGGCCG*GTAGCGAGTGATAGCGAGTGAAAGACGATCGCTATCGTGCCGTAACG AGGAAATGTGAGGTTGGGGG. EaeI digestion of the DNA fragment was used to identify the clones harboring mutations of upstream Adh1 G4 motif. Positive clones were confirmed by sequencing the regions around the *adh1^+^* locus.

### ChIP using the BG4 antibody

BG4 antibody was purified as previously described (24). 10 mL exponentially growing *S. pombe* cells (2 x 10^6^ cells/mL) were crosslinked in 1% formaldehyde for 10 min and the reactions were quenched with 125 mM final concentration of glycine. Cells were harvested and lysed in ChIP lysis buffer (50 mM Hepes/KOH (pH 7.5), 140 mM NaCl, 1 mM EDTA, 1% Triton X-100, and 0.1% Na-deoxycholate) using glass beads in a FastPrep-24^TM^ benchtop cell homogenizer (MP Biomedicals) at 4 °C. Chromatin was isolated and sheared using a Covaris E220 system to an average size of 100 to 600 bp. After RNase A treatment for 30 min, chromatin was incubated with 0.8 µg BG4 antibody in ChIP lysis buffer containing 1% milk in a thermomixer shaking at 1200 rpm at 16 °C for 1 h. 8 µg anti-FLAG antibody (Sigma Aldrich #F3165) was bound to 10 µL protein-G magnetic beads (Pierce™ ThermoFisher) by shaking in a thermomixer at 1200 rpm at 16 °C for 2 h. The anti-FLAG-bound magnetic beads were washed with ChIP lysis buffer and incubated with chromatin-BG4 complex in a thermomixer shaking at 1200 rpm at 16 °C for 2 h. Beads were washed six times with ChIP lysis buffer and twice with wash buffer (100 mM KCl, 0.1% (w/v) Tween-20, 10 mM Tris HCl pH 7.5). The immunoprecipitated DNA was eluted with 75 µL TE buffer and 0.5 mg/mL Proteinase K by incubating at 37 °C for 1 h followed by 65 °C for 2 h shaking in the thermomixer. Finally, the eluates were purified using ChIP DNA Clean and Concentrate kit (ZYMO Research). qPCR was performed using primer pairs amplifying the upstream *adh1* G4 or *ade6* (non-G4 control) regions.

### ChIP using Histone H3 tri methyl lysine 4 (H3K4me3) antibody

Cellular growth, crosslinking and chromatin isolation were performed as described in BG4-ChIP above. H3K4me3 antibody (Abcam) was incubated with chromatin overnight at 4 °C. Next, 40 µL protein-G magnetic beads (Pierce™ ThermoFisher) was added to the chromatin-antibody mixture and incubated at 4 °C for 4.5h. The magnetic beads were sequentially washed with SDS buffer (50 mM HEPES-KOH, pH7.5, 140 mM NaCl, 1 mM EDTA, 0.025% SDS), High Salt Wash Buffer (50 mM Hepes-KOH, pH 7.5, 1 M NaCl, 1mM EDTA), T/L Wash Buffer (20 mM Tris-HCl, pH 7.5, 0.25 M LiCl, 1 mM EDTA, 0.5% Sodium Deoxycholate, 0.5% IGEPAL-CA630) and T/E Buffer (20 mM Tris-HCl, pH 8.0, 0.1 mM EDTA). The immunoprecipitated DNA was eluted with 145 µL Elution buffer (20 mM Tris-HCl, pH 8.0, 0.1 mM EDTA, 1% SDS) by heating at 65 °C for 2 mins. Reverse crosslinking was performed overnight at 65 °C. Subsequently, 4 µL Proteinase K (20mg/mL) was added and incubated at 50 °C for 2h. Finally, DNA was purified using ChIP DNA Clean and Concentrate kit (ZYMO Research). qPCR was performed using a set of primer pairs that span the Adh1 G4 site up to the ATG start codon of *adh1* gene or *act1* gene (control).

### Dot blot assay for G4 structure detection

0.25 µM DNA oligonucleotides were folded by heating at 95 °C in 100 mM KCl or water and allowed to cool down to room temperature. 200 µL of the DNA oligonucleotides was loaded on Hybond-N+ membrane (GE Healthcare) using a Bio-Dot Microfiltration Apparatus (Bio-Rad) and gentle vacuum. After blocking with 5% milk in intracellular salt solution (25 mM Hepes, pH 7.5, 10.5 mM NaCl, 110 mM KCl and 1 mM MgCl_2_), the membrane was incubated with the BG4 antibody (0.4 µg/mL) overnight at room temperature and washed twice for 15 min with wash buffer (10 mM Tris, pH 7.4, 100 mM KCl, 0.1 % (vol/vol) Tween 20). The membrane was incubated with mouse anti-FLAG antibody (Sigma-Aldrich, #F3165, 1:5000 dilution) for 2 h at room temperature and washed twice for 15 min with the wash buffer. Finally, the membrane was incubated with Goat anti-mouse IgG (H+L) antibody-HRP (Thermo Fisher Scientific) (1:5000 dilution) for 1 hour at room temperature and dots were visualized using chemiluminescent reagents (Pierce SuperSignal™ West Pico PLUS Chemiluminescent Substrate).

### Real-time quantitative PCR (RT-qPCR)

Total RNA was isolated from exponentially growing *S. pombe* cells using NucleoSpin RNA kit (Macherey Nagel). After checking the quality and integrity of the isolated RNA using absorbance (260/280 nm) and agarose gel electrophoresis, cDNA was made using UltraScript 2.0 cDNA synthesis kit (PCR Biosystems). To check for genomic DNA contamination, a control without the reverse transcriptase was included. Primers amplifying coding regions of *adh1^+^* and *ade6^+^*genes were used to quantify *adh1^+^* and *ade6^+^*gene expression, respectively. Levels of *act1^+^* and *tdh1^+^* were used as reference genes for normalization (Supplementary Table 2).

### Bulk RNA sequencing

Total RNA was isolated from ADH1 G4 WT, ADH1 G4-TATA MUT and ADH1 G4 MUT strains grown in Yeast Extract Supplement (YES) media. Messenger RNA was then purified from the total RNA using poly-T oligo-attached magnetic beads. Following fragmentation, the first strand of cDNA was synthesized using random hexamer primers. This was followed by the synthesis of the second strand cDNA, utilizing either dUTP for the creation of a directional library or dTTP for a non-directional library. The constructed library was assessed using Qubit and real-time PCR for quantification, as well as a bioanalyzer to determine the size distribution. Quantified libraries were subsequently pooled and sequenced on high-throughput Illumina sequencing platforms at Novogene sequencing Europe. The sequenced data, referred to as Raw Data or Raw Reads, were obtained in FASTQ format files. The sequencing reads were filtered to remove those with adapter contamination, low-quality nucleotides, and uncertain nucleotides. The filtered reads were aligned to the yeast reference genome (972h-Genome assembly ASM294v2) using HISAT2. Mapped regions were classified as exonic, intronic, or intergenic and were annotated accordingly. Gene expression levels were estimated based on the abundance of transcripts that mapped to the genome or exons, using the FPKM (Fragments Per Kilobase of transcript sequence per Millions base pairs sequenced) method. The correlation of gene expression levels between samples was quantified using the Pearson correlation coefficient. Principal component analysis (PCA) was performed to evaluate intergroup differences and intragroup sample duplication. Differential gene expression analysis was conducted using DESeq2, with thresholds set at |log2(FoldChange)| ≥ 1 and padj ≤ 0.05. For functional analyses, GO enrichment was performed using GSEA, and significantly affected pathways were selected based on a padj value of < 0.05.

### Metabolomic analysis

A cell number of 2.3 x 10^8^ logarithmic phase *S. pombe* cells grown in YES were washed three times in cold Phosphate Buffered Saline (PBS). Cell pellets were stored at −80 °C. Metabolic profiling by gas chromatography – mass spectrometry (GC – MS) was performed at the Swedish Metabolomic Centre (SMC) at Umeå university, Sweden. Sample preparation was performed according to Gullberg et al (78). In detail, metabolites were extracted from the yeast cells using 1200 µL of extraction buffer (90/10 v/v methanol: water) together with internal standards (L-proline-13C5, alpha-ketoglutarate-13C4, myristic acid-13C3, cholesterol-D7, succinic acid-D4, salicylic acid-D6, L-glutamic acid-13C5,15N, putrescine-D4, hexadecanoic acid-13C4, D-glucose-13C6, D-sucrose-13C12) by shaking with one tungsten bead in a mixer mill at 30 Hz for 2 minutes. The bead was removed, and the samples were centrifuged at +4 °C, 14 000 rpm (18,620 g) for 10 minutes. 25 µl of the supernatant was transferred to micro vials and evaporated to dryness in a speed-vac concentrator. Solvents were evaporated and the samples were stored at −80 °C until analysis. A small aliquot of the remaining supernatants was pooled and used to create quality control (QC) samples. The samples were analyzed according to a randomized run order.

Derivatization and GC-MS analysis were performed as described previously (78). 1 μL of the derivatized sample was injected in splitless mode by a L-PAL3 autosampler (CTC Analytics AG, Switzerland) into an Agilent 7890B gas chromatograph equipped with a 10 m x 0.18 mm fused silica capillary column with a chemically bonded 0.18 μm Rxi-5 Sil MS stationary phase (Restek Corporation, U.S.) The injector temperature was 270 °C, the purge flow rate was 20 mL min^-1^ and the purge was turned on after 60 seconds. The gas flow rate through the column was 1 mL min-1, the column temperature was held at 70 °C for 2 minutes, then increased by 40 °C min-1 to 320 °C, and held there for 2 minutes. The column effluent was introduced into the ion source of a Pegasus BT time-of-flight mass spectrometer, GC/TOFMS (Leco Corp., St Joseph, MI, USA). The transfer line and the ion source temperatures were 250 °C and 200 °C, respectively. Ions were generated by a 70 eV electron beam at an ionization current of 2.0 mA, and 30 spectra s-1 were recorded in the mass range m/z 50 - 800. The acceleration voltage was turned on after a solvent delay of 150 seconds. The detector voltage was 1800-2300 V.

All non-processed MS-files from the metabolic analysis were exported from the ChromaTOF software in NetCDF format to MATLAB R2021a (Mathworks, Natick, MA, USA), where all data pre-treatment procedures, such as base-line correction, chromatogram alignment, data compression and Multivariate Curve Resolution were performed. The extracted mass spectra were identified by comparisons of their retention index and mass spectra with libraries of retention time indices and mass spectra (79). Mass spectra and retention index comparison was performed using NIST MS 2.2 software. The annotation of mass spectra was based on reverse and forward searches in the library. Masses and ratio between masses indicative of a derivatized metabolites were especially notified. The mass spectrum with the highest probability indicative of a metabolite and the retention index between the sample and library for the suggested metabolite was ± 5 (usually less than 3) the deconvoluted “peak” was annotated as an identification of a metabolite.

### qPCR stop assay

The qPCR stop assay was performed as described previously (71). Briefly, reactions were performed with primer pairs targeting the upstream Adh1 G4 or Ade6 (non-G4 control) (Supplementary Table 2) regions and contained 0.3 µM primer, 33 ng *S. pombe* genomic DNA, and 1x SyGreen mix (PCR Biosystems) in the presence of either 0.65% (v/v) DMSO, 25 mM KCl, or 25 mM KCl and 0.5 µM (Phenanthroline dihydrochloride-3,6-diamino-9-[2-(2-carboxyethyl)-3-hydroxy-4-methyl-1H-pyrrol-1-yl]-2,7-dimethyl-9H-fluorene-2,7-dicarboxylic acid dihydrochloride) (PhenDC3) mixture. The PCR program was 95 °C for 5 min (1 cycle) followed by a 2-step reaction of 95 °C for 10 s and 60 °C for 20 s (33 cycles) in one-point acquisition mode. ΔΔ*Cq* values were determined to express the relative DNA amplification.

### Determination of doubling times of *S. pombe* strains

2 x 10^5^ cells/mL *S. pombe* cells in logarithmic phase were grown in YES media at 30 °C and 140 rpm for 16 h. Cells were counted under the microscope using a Bürker chamber. The experiments were performed three times, independently. Doubling times were calculated using the following formula: Doubling time = *t* / log_2_ (*x /x_0_*), where *t* is time in hours; *x* is counted cell number at 16 h and *x_0_* is counted cell number at 0 h.

### Native gel electrophoresis for G4 structure determination

0.25 µM ADH1 WT, ADH1 5_G MUT or ADH1 4_G MUT oligonucleotides were labeled using T4 polynucleotide kinase (PNK) (Thermo Fisher Scientific) and γ-^32^P-ATP at 37 °C for 75 min. T4 PNK was inactivated by addition of 25 mM EDTA and heating at 78 °C for 1 min. Radiolabeled oligonucleotides were purified on a G50 column (GE Healthcare). 50 nM oligonucleotides were folded in folding buffer (10 mM Tris pH 7.5, 100 mM KCl) by heating at 95 °C for 5 min and allowing the oligonucleotides to cool down to room temperature. The oligonucleotides were analyzed on a 15% native polyacrylamide gel containing 50 mM KCl at 100 V for 80 min. Bands were visualized using Amersham Typhoon Scanner (GE Healthcare) and analyzed by the ImageQuant 5.2 software (GE Healthcare).

### Circular dichroism (CD) spectroscopy

50 µM ADH1 WT, ADH1 5_G MUT or ADH1 4_G MUT oligonucleotides were heated in water or folding buffer (10 mM Tris pH 7.5, 100 mM KCl) at 95 °C for 5 min and allowed to fold at room temperature for 3 h. Folded oligonucleotides were diluted to 5 µM. For the experiments with PhenDC3, 10 µM of the ligand or equivalent volume of DMSO was added. CD spectra were recorded between 225 and 350 nm at 25 °C using a JASCO-1700 spectrometer with a Peltier temperature control in a quartz cuvette (0.1 cm path length). Each spectrum was the accumulation of three measurements. Baseline corrections were performed using folding buffer containing either 1.25% (v/v) DMSO or 10 µM PhenDC3. All data were normalized to molar ellipticity using the formula: [*θ*] = *m°*× *M* /10 × *L* × *C*

Where *m◦* is CD signal in millidegrees; *M* is molecular weight of oligonucleotides in g/mol; *L* is path length of cell in cm; *C* is concentration of oligonucleotides in g/l.

### 1H NMR measurements

120 µM ADH1 WT, ADH1 5_G MUT, and ADH1 4_G MUT oligonucleotides were folded in 100 mM KCl by heating to 95 °C and slowly cooling to room temperature. 200 µL of the oligonucleotide solution in presence of 20 mM Tris (pH 7.5) and 10% D_2_O were loaded into 3 mm NMR tube. Spectra were recorded from 25 °C to 45 °C on a Bruker 850 MHz Avance III HD spectrometer equipped with a 5 mm TCI cryoprobe. Excitation sculpting was used, and 256 scans were recorded. Topspin 4.1.3 (Bruker Biospin, Germany) was used for data processing.

## RESULTS

### G4 CRISPR-Cas9 - an effective tool for generation of *S. pombe* strains harboring mutations within Adh1 G4 motif

We employed the CRISPR-Cas9 system to introduce multiple G-to-T point mutations simultaneously in an endogenous G4 motif of *S. pombe* cells. Based on the studies performed in the Allshire lab (62), which successfully generated single point mutations in *S. pombe* genome, we designed a CRISPR-Cas9 mutational approach to simultaneously target multiple G residues within the G4 motif (Figure 1A and Supplementary Figure 1). This approach, hereafter referred to G4 CRISPR-Cas9, was applied to a previously identified G4 site (marked as G435 in Figure 1B), located upstream of the *adh1^+^* gene on the antisense/template strand, henceforth referred to as the Adh1 G4 motif. The Adh1 G4 motif has been shown to cause increased DNA polymerase occupancy, suggesting slower replication fork progression in this region (Figure 1B) (67). To modify the Adh1 G4 motif, we designed two sgRNAs targeting distinct positions within the motif and successfully integrated them into a pLSB-NAT plasmid (Figure 1A and Supplementary Figures 2A, 2B, and 2C). Next, we performed recombinational PCR to create two homologous recombination (HR) templates (HR1 and HR2) (Supplementary Figure 2D), designed to match either the ADH1 5_G MUT or ADH1 4_G MUT sequences. The ADH1 5_G MUT oligonucleotide sequence contains four G-to-T mutations within the G4 motif, as well as a G-to-C substitution introduced to disrupt the PAM sequence (AGG to ACG) necessary for CRISPR-Cas9 editing, while the ADH1 4_G MUT oligonucleotide sequence carried four G-to-T mutations, inherently disrupting the PAM sequence (CGG to CGT) (Figure 2A). Haploid *S. pombe* cells were transformed with sgRNA-containing plasmids and HR templates. Successful genomic recombination was confirmed in 20% of screened clones (Supplementary Figures 2E and 2F), and sequencing verified the integration of HR1 and HR2, resulting in two mutant strains.

**Figure 1.**
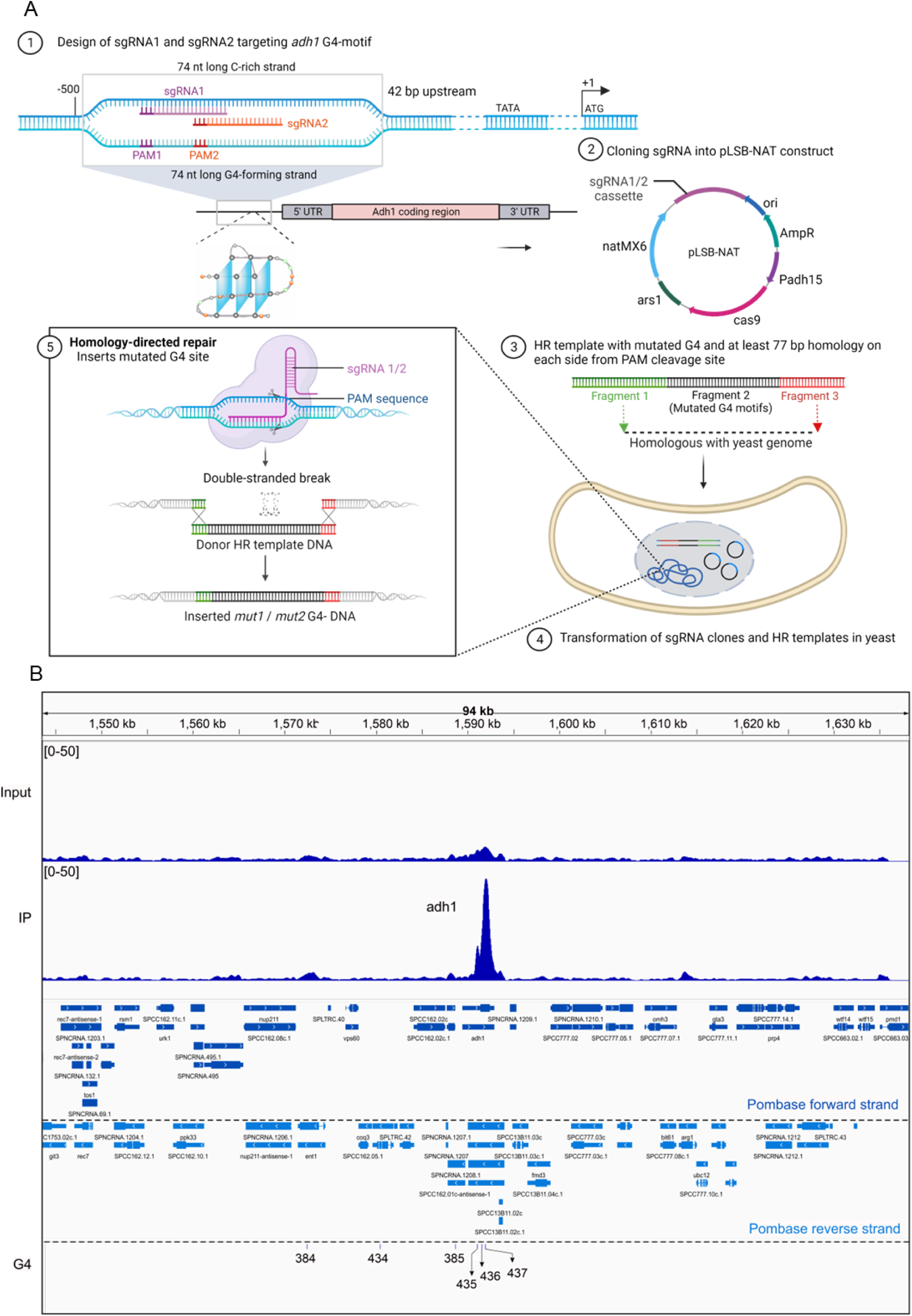
G4 CRISPR-Cas9 strategy to mutate the G4 motif at *adh1^+^* promoter in *S. pombe*. (**A**) Schematics illustrating important steps for mutating Adh1 G4 motif using G4 CRISPR-Cas9 system in *S. pombe*. In short, sgRNAs targeting simultaneously different sites within the G4 motif were designed and Golden Gate Cloning that allows scarless cloning in a single step was used to insert the sgRNAs into a plasmid (pLSB-NAT) containing a Green Fluorescent Protein (GFP) gene for simple selection of *E. coli* transformants with sgRNA-containing plasmids. Homologous recombination (HR) templates were designed to carry the desired G4 motif mutations, which were introduced together with the pLSB-NAT into haploid cells through transformation and incorporated into the genome by homology-directed repair. Created by Biorender.com (**B**) Genome browser track showing Cdc20 (catalytic subunit of DNA polymerase ε) ChIP-seq datasets (European Nucleotide Archive, accession number: PRJEB37862) performed in *S. pombe* (67). The Cdc20 enrichment profiles across *adh1^+^*genomic region is shown, where the y-axis represents the normalized read density, reflecting the relative abundance of sequencing reads at each genomic position. The normalized read density values have been scaled to a range of 0 to 50 to facilitate visual comparison between the input and Cdc20 immunoprecipitant (IP) samples. The labels below the tracks show the Pombase forward and reverse strands; and the specific position of bioinformatically predicted G4s in chromosome III. The Adh1 G4 motif is denoted as G435 in the G4 track and identified using the algorithm (G_≥3_ N_1–25_)_3_ G_≥3,_ where N can be 1-25 nucleotides (9).

**Figure 2.**
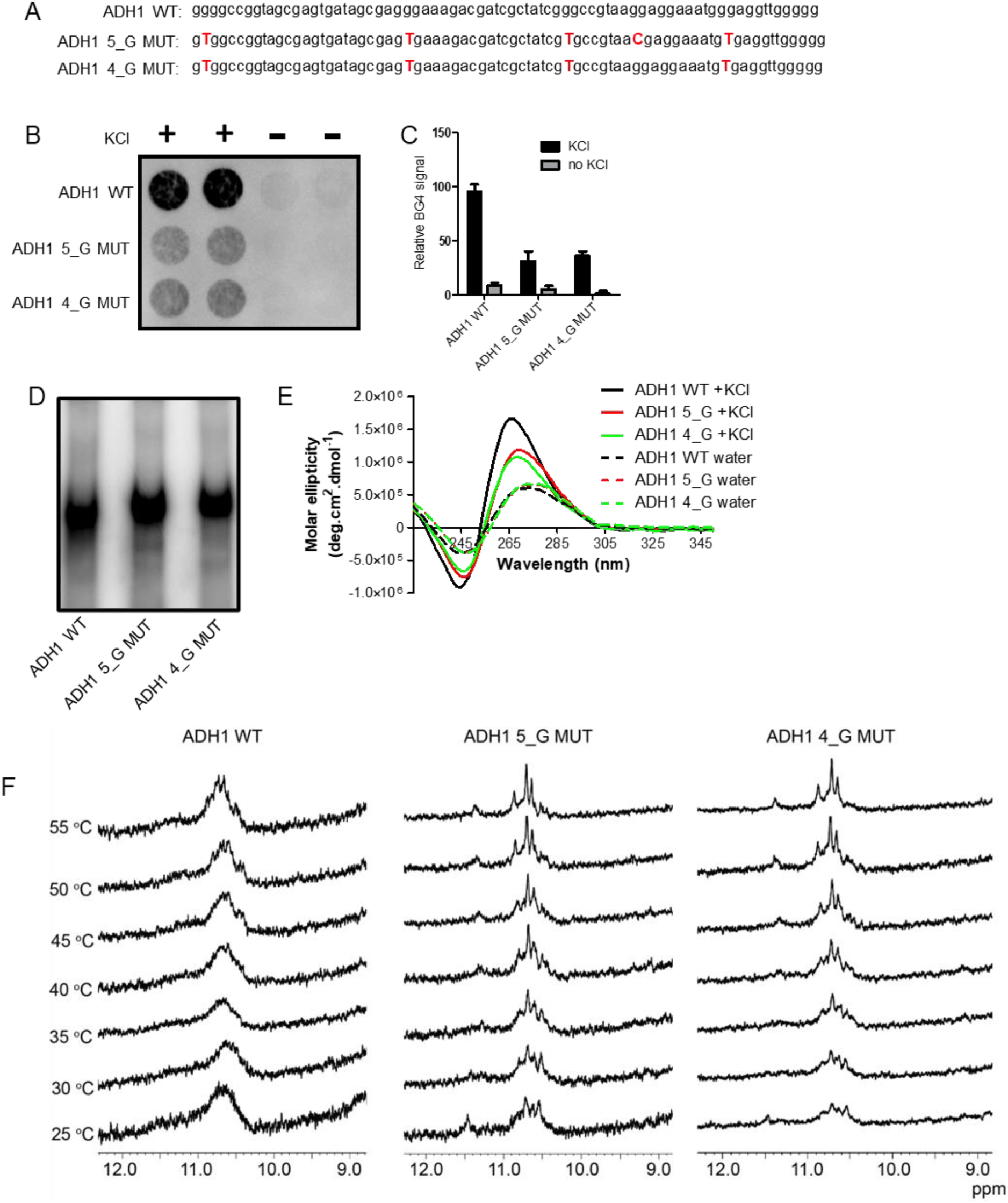
*In vitro* validation of G4 structure properties of oligonucleotides having native or mutated Adh1 G4 motif. (**A**) Sequences of oligonucleotides with native (ADH1 WT) or mutated (ADH1 5_G MUT and ADH1 4_G MUT) Adh1 G4 motif. The red, bold, and capitalized T and C represent thymine and cytosine residues intended to modify native G4 structure formation. The presence of cytosine (red capital C) instead of guanine in ADH1 5_G MUT distinguishes this oligonucleotide from ADH1 4_G MUT. (**B**) BG4 dot blot assay to probe G4 structure formation. ADH1 WT, ADH1 5_G MUT, and ADH1 4_G MUT oligonucleotides were folded with or without 100 mM KCl. G4 structure formation was detected using BG4 antibody. Two replicates are shown for each oligonucleotide and condition. (**C**) Quantification of BG4 signals of the dot blot from B. The graph shows the average values of two experiments. Error bars represent the standard deviation. (**D**) Native gel electrophoresis to examine the molecularity of ADH1 WT, ADH1 5_G MUT and ADH1 4_G MUT oligonucleotides. Radiolabeled oligonucleotides were folded in 100 mM KCl and run on a 10% native polyacrylamide gel. (**E**) CD spectra of ADH1 WT, ADH1 5_G MUT and ADH1 4_G MUT oligonucleotides. The oligonucleotides were folded in water or 100 mM KCl. The CD spectra were recorded between 225 and 350 nm. OriginLab 2020 was used to smoothen the curves. (**F**) ^1^H NMR spectroscopy of ADH1 WT, ADH1 5_G MUT and ADH1 4_G MUT DNA oligonucleotides at different temperatures. The oligonucleotides were folded with KCl. NMR spectra were recorded at different temperatures on a Bruker 850 MHz Avance III HD spectrometer equipped with a 5 mm TCI cryoprobe and analyzed by Topspin 4.1.3 software.

One strain contained mutations exclusively within the Adh1 G4 motif, hereafter referred to as the ADH1 G4 MUT strain. The second strain exhibited two additional point mutations, likely caused by the sgRNA plasmid (see Discussion). These mutations were detected in the TATA-box, located 322 and 324 base pairs downstream of the Adh1 G4 motif. Although the TATA-box mutations were unintended, we included this strain as a control, hereafter referred as the, ADH1 G4-TATA MUT strain. Including this strain provided a valuable reference for understanding Adh1 gene regulation, as TATA-box mutations significantly compromise transcription (80), and allowed us to isolate the specific effects of the G4 mutation in ADH1 G4 MUT through direct comparison.

### Mutations alter G4 folding propensity *in vitro*

Before investigating the *S. pombe* strains, we characterized the G4 folding properties of the mutated and wild-type ADH1 G4 motifs. Using the G4 Killer web application (version 7/21/2021) (81), we assessed whether the designed sequences had a reduced propensity to fold into G4 structures. This tool, based on the G4 Hunter algorithm, predicts G4 formation in mutated motifs (7). For this analysis, we used oligonucleotides corresponding to the ADH1 WT, ADH1 5_G MUT, and ADH1 4_G MUT sequences, matching the designs used in strain construction (Figure 2A). Compared to the ADH1 WT sequence, which got the G4 Hunter score of 1.01, the scores for ADH1 5_G MUT and ADH1 4_G MUT oligonucleotide sequences were reduced to 0.53 and 0.58, respectively, predicting that the mutations reduced the propensity of these sequences to fold into G4 structures.

To assess whether the mutations affected G4 structure formation *in vitro*, we folded the ADH1 WT, ADH1 5_G MUT, and ADH1 4_G MUT oligonucleotides in the presence or absence of KCl, as potassium ions promote G4 folding (4). We then performed a dot blot assay using BG4 antibody, which recognizes different types of G4 structures (Figures 2B). The ADH1 WT oligonucleotide folded in KCl showed a pronounced BG4 antibody signal, while BG4 signals for the ADH1 5_G MUT and ADH1 4_G MUT oligonucleotides folded in KCl were more than three times lower (Figure 2C). Importantly, when folded without KCl, all oligonucleotides exhibited negligible BG4 signal (Figure 2B, 2C), showing that G4 folding is KCl-dependent. These results indicate that the G residue mutations in the oligonucleotides partially compromise G4 structure formation, and that a single nucleotide mutation (G to C) between ADH1 5G_MUT and ADH1 4G_MUT does not significantly affect recognition by BG4.

Next, we used native polyacrylamide gel electrophoresis (PAGE) to evaluate whether oligonucleotide sequence modifications affected the molecularity of the oligonucleotides in forming intra- or intermolecular G4 structures. On a native gel, intramolecular G4 structures, which form within a single DNA molecule, migrate faster due to their compact nature compared to ssDNA. In contrast, intermolecular G4 structures formed by two or more DNA molecules, migrate more slowly because of their higher molecular weight (82). None of the three oligonucleotides folded into intermolecular G4 structures (Figure 2D). The ADH1 WT oligonucleotide folded in KCl migrated faster than the mutated counterparts (ADH1 5_G MUT and ADH1 4_G MUT), suggesting that it forms more compact intramolecular G4 structures (Figure 2D). To validate these results and determine the G4 topologies of the formed structures, we performed circular dichroism (CD) spectroscopy (83). The ADH1 WT oligonucleotide folded in KCl displayed a positive peak at 266 nm and a negative peak at 245 nm, consistent with the formation of a parallel G4 structure (Figure 2E) (83). In contrast, the mutated oligonucleotides showed significant hypochromic effects and slightly red-shifted positive peaks (∼ 269 nm) (Figure 2E), indicating that the mutations partially compromised or altered G4 folding in both sequences. To investigate further, we titrated PhenDC3 (1_DNA_:2_PhenDC3_) into these G4-forming sequences followed by acquisition of CD spectra. PhenDC3 demonstrates higher selectivity for G4 structures compared to single-stranded DNA (84). Also, it can induce G4 conformational changes (85,86). In the presence of PhenDC3, a slight broadening of the spectra was observed with all three oligonucleotides, suggesting interactions between PhenDC3 and the three oligonucleotides (Supplementary Figure 4) (85,86). Additionally, when folded in water, none of the three oligonucleotides showed typical G4 spectra, supporting the dot blot assays results (Figure 2B and 2C).

### Reduced structural dynamics in mutated sequences by NMR spectroscopy

While CD spectroscopy provides valuable insights into G4 topologies, its sensitivity to detect subtle structural differences is limited. To better understand the structural variations in these oligonucleotides, we examined their 1D ^1^H nuclear magnetic resonance (NMR) spectra in the presence or absence of KCl (Figure 2F and Supplementary Figure 5). The spectra of all three sequences were influenced by KCl, showing characteristic G4 signals around 10 and 12 ppm, corresponding to the imino protons of the Gs involved in Hoogsteen hydrogen bonding (87). Notably, the ADH1 5_G MUT and ADH1 4_G MUT oligonucleotides showed multiple imino proton signals with sharper linewidths, in contrast to the broad envelope of imino proton resonances observed in the ADH1 WT spectra. The broad imino signals in the ADH1 WT indicate a heterogeneous ensemble of interconverting G4 structures, while the sharper signals in the mutated sequences suggest that the G-to-T mutations restrict structural dynamics, resulting in distinct G4 variants. To further probe these differences, NMR spectra were recorded at various temperatures. The imino proton resonances (10–12 ppm) in the mutated sequences remained sharper than those in the wild-type, indicating that the G4 structures in the mutated oligonucleotides adopt more well-defined conformations. Despite having an additional G-to-C mutation in ADH1 5_G MUT, both mutants did not exhibit significant differences in terms of structural dynamics. In contrast, the ADH1 WT G4 maintained a broad envelope of imino signals even at 45 °C, reflecting greater structural heterogeneity and dynamic interconversion. These data suggest that the Adh1 G4 can adopt multiple G4 conformations, similar to what has been observed with other G4 sequences, for example the human telomeric G4 DNA (88,89).

While the introduced mutations compromised and partially altered G4 formation and dynamics, they did not completely disrupt the folding potential. This highlights the ability of the mutated sequences to form G4 structures and underscores the differences in structural plasticity between G4s formed by the ADH1 WT and the mutated oligonucleotides.

### G4 motif upstream of *adh1^+^* gene forms G4 structure in the genome

To determine whether the ADH1 WT G4 motif and the mutated sequences form stable G4 structures within genomic DNA, we isolated genomic DNA from ADH1 G4 WT, ADH1 G4-TATA MUT, and ADH1 G4 MUT *S. pombe* strains and performed a qPCR stop assay (Figure 3A) (71). In this assay, G4 structure formation in the isolated genomic DNA inhibits DNA synthesis, resulting in reduced amplification. This *in vitro* assay evaluates the efficiency of Taq DNA polymerase in synthesizing the genomic region of interest in the presence or absence of a G4 motif and can be combined with small molecules that stabilize or destabilize G4 structures (67,71,90). We designed primer pairs that specifically annealed to the Adh1 G4 region or a non-G4 control region (Ade6) (Supplementary table 2). Genomic DNA from the ADH1 G4 WT strain showed significantly reduced amplification of the Adh1 G4 region compared to DNA from ADH1 G4-TATA MUT and ADH1 G4 MUT strains, which exhibited 3- and 1.5-fold higher amplification, respectively (Figure 3B).

**Figure 3.**
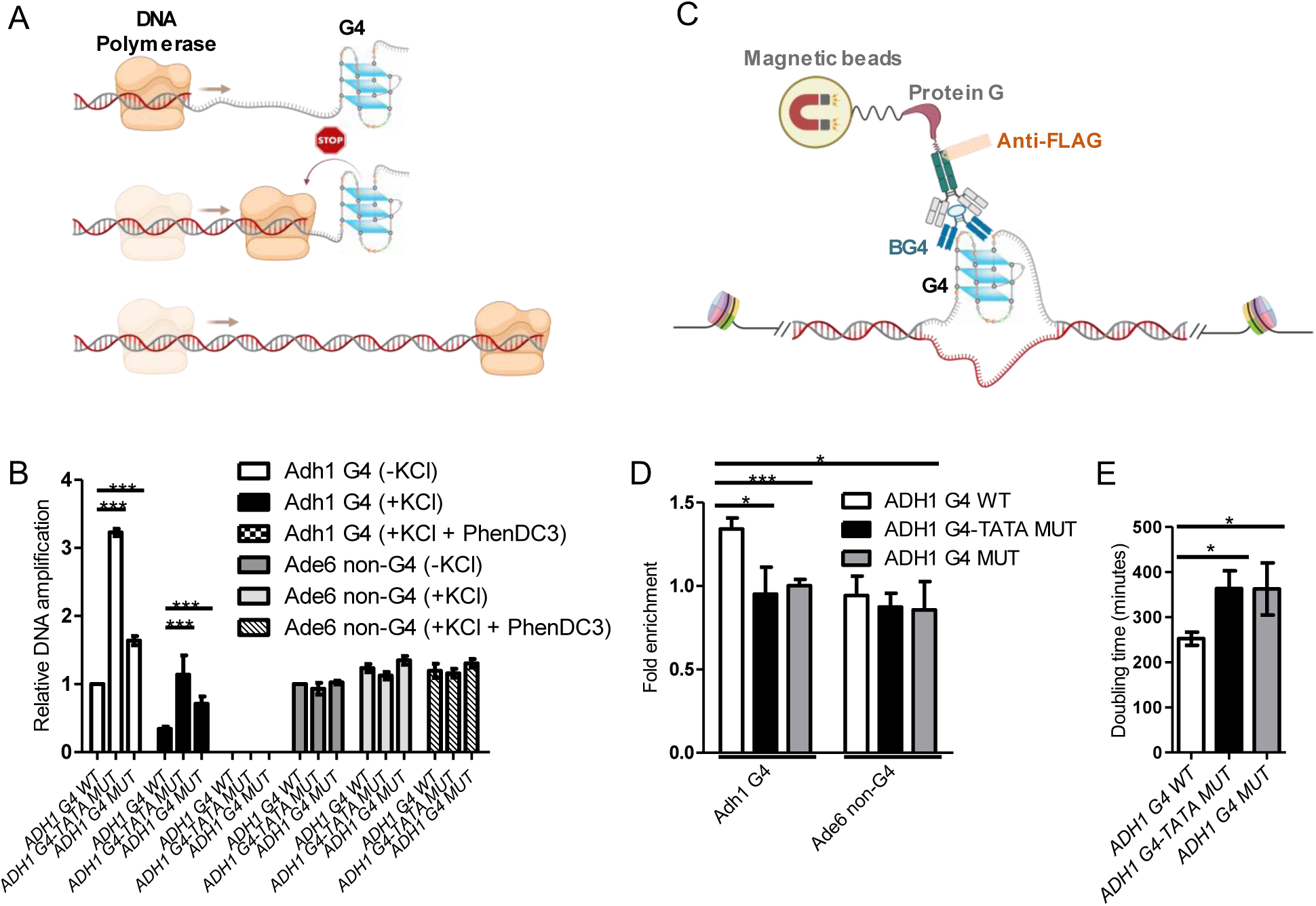
Functional analysis of Adh1 G4 motif in *S. pombe*. (**A**) Schematics showing the principles for the qPCR stop assay. Created by Biorender.com (**B**) Genomic DNA from *S. pombe* strains harboring ADH1 WT or mutated (ADH1 5_G MUT or ADH1 4_G MUT) Adh1 G4 motifs were used as templates in qPCR stop assay performed in the absence of KCl or in the presence of KCl, with or without PhenDC3. The primer pairs were designed to flank the G4 or non-G4 (*ade6*+) sites. The graph shows the average values of three experiments. Error bars represent the standard deviation. * *P* < 0.05 and *** *P* < 0.001 according to two-sample *t*-test. (**C**) Schematics showing the principles for BG4-ChIP. Created by Biorender.com (**D**) DNA from chromatin isolated from *S. pombe* strains harboring ADH1 G4 WT or mutated (ADH1 G4-TATA MUT or ADH1 G4 MUT) Adh1 G4 motifs were immunoprecipitated using BG4 antibody. The amount of immunoprecipitated DNA was analyzed by qPCR using primer pairs that flank the G4 or non-G4 (*ade6*+) sites. Input C_q_ value of each sample was used in normalization. The graph shows the average values of four experiments. Error bars represent the standard deviation. * *P* < 0.05 and *** *P* < 0.001 according to two-sample *t*-test. (**E**) Growth of *S. pombe* strains harboring ADH1 G4 WT or mutated (ADH1 G4-TATA MUT or ADH1 G4 MUT) Adh1 G4 motifs. The graph shows the average values of the doubling times of three experiments. Error bars represent the standard deviation. * *P* < 0.05 according to two-sample *t*-test.

Consistent with these findings, the addition of KCl further reduced DNA amplification across all sequences, likely due to enhanced G4 formation, while maintaining the observed trend (Figure 3B). These results suggest that the mutated regions can form G4 structures in the presence of KCl, but these structures are less stable than those in the ADH1 G4 WT region. When PhenDC3 was combined with KCl, DNA amplification of the Adh1 G4 was completely inhibited in DNA isolated from all strains, showing that PhenDC3 and KCl together significantly enhanced G4 stability of all three sequences. In contrast, amplification of the non-G4 Ade6 control region remained unchanged under all conditions and across all strains (Figure 3B), confirming that the observed effects were specific to the Adh1 G4 site in the isolated genomes.

Next, we determined the G4 structure formation in the Adh1 G4 region within chromatin using ChIP with the G4-binding BG4 antibody, followed by qPCR analysis in the three strains (Figure 3C). The ChIP-qPCR experiments were performed with two primer pairs that either targeted the Adh1 G4 region or the Ade6 non-G4 control region (Figure 3D). Based on our *in vitro* findings that point mutations in the ADH1 5_G MUT and ADH1 4_G MUT sequences compromised the potential formation of stable G4 structures, we hypothesized that G4 levels at the Adh1 G4 region (as detected by BG4 antibody) would be higher in the ADH1 G4 WT strain compared to the mutated strains. Indeed, BG4 occupancy at the Adh1 G4 region was significantly enriched in the ADH1 G4 WT strain compared to both the mutated strains, as well as the negative control, the Ade6 non-G4 region (Figure 3D). These results confirm the *in vivo* formation of G4 structures at the Adh1 G4 region in ADH1 G4 WT cells and demonstrate that G4 formation at this site is compromised in the mutated strains, aligning with our observations from the BG4 dot blot and other *in vitro* experiments (Figures 2 and 3B).

Furthermore, we evaluated the biological impact of these point mutations on cellular growth. *S. pombe adh1*Δ cells are viable, but they exhibit slower growth compared to WT cells (91). Similarly, both the ADH1 G4-TATA MUT and ADH1 G4 MUT strains showed significantly impaired growth rates compared to ADH1 G4 WT cells, suggesting that the Adh1 G4 is important for optimal cellular growth (Figures 3E).

### Adh1 G4 motif positively regulates *adh1^+^* transcription

Since mutations in the Adh1 G4 in ADH1 G4-TATA MUT and ADH1 G4 MUT cells affected cellular growth, we explored whether these effects are due to dysregulation of *adh1^+^* gene expression. As described earlier in this manuscript, in addition to the point mutations in the Adh1 G4 motif, the ADH1 G4-TATA MUT cells also contained two mutations (T to A and A to T at positions 106 and 104 base pairs) within the TATA box upstream of the *adh1^+^* start codon (Supplementary Figure 3). Previous studies have demonstrated that the TATA elements upstream of the *adh1^+^* start site are essential for proper gene expression levels (80). Therefore, the ADH1 G4-TATA MUT cells were used as a control in our gene expression analyses. mRNA isolated from ADH1 G4 WT, ADH1 G4-TATA MUT, and ADH1 G4 MUT cells were reverse transcribed to cDNA and quantified using primer pairs that amplified the coding region of *adh1*^+^ or *ade6*^+^ (non-G4 control) genes. As expected from earlier findings on the *adh1^+^* TATA box (80), ADH1 G4-TATA MUT cells showed a pronounced reduction in *adh1^+^* mRNA levels, with a more than four-fold decrease compared to ADH1 G4 WT cells. While ADH1 G4 MUT cells also exhibited lower *adh1^+^* expression, the decrease was less pronounced than in ADH1 G4-TATA MUT cells (Figure 4A). Thus, the reduced *adh1^+^*expression in ADH1 G4-TATA MUT cells cannot be conclusively attributed solely to TATA box mutations. Mutations in the Adh1 G4 motif may also contribute to this reduction, as the ADH1 5_G MUT and ADH1 4_G MUT sequences exhibited similar behaviors in prior *in vitro* and *in vivo* G4 characterizations. However, further studies are necessary to disentangle the relative contributions of TATA box and G4 mutations to the observed effects on *adh1^+^*expression. Together, these findings demonstrate that the Adh1 G4 motif plays a positive (92) regulatory role in *adh1^+^* expression, aligning with recent studies by the Balasubramanian and Di Antonio labs on the G4 motif in the human *cMYC* promoter (55,57).

**Figure 4.**
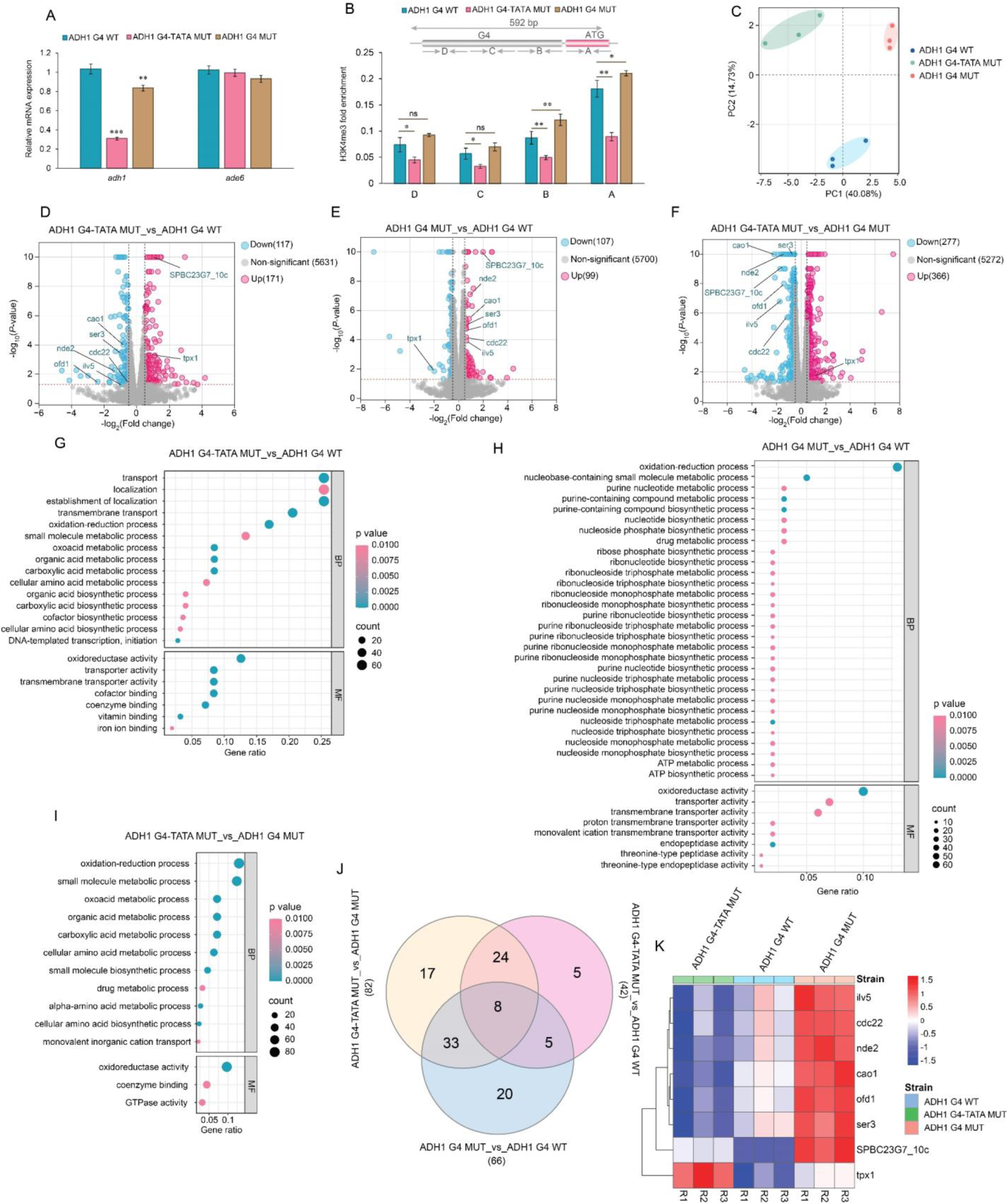
Mutation of Adh1 G4 affects *adh1* levels and alters global cellular transcriptome (**A**) Realtime qPCR for *adh1^+^* gene expression. Total RNA was isolated from *S. pombe* strains harboring ADH1 G4 WT or mutated (ADH1 G4-TATA MUT or ADH1 G4 MUT) Adh1 G4 motifs and reverse transcribed to cDNA using a combination of random hexamers and anchored oligo (dT). Primer pairs amplifying the *adh1^+^* and *ade6^+^* (non-G4) coding regions were used. *act1^+^* and *tdh1^+^* were used as reference genes for normalization. The graph shows the average values of three experiments. Error bars represent the standard deviation. ** *P* < 0.01 and *** *P* < 0.001 according to two-sample t-test. (**B**) H3K4me3 - ChIP to probe *adh1^+^* locus. DNA from chromatin isolated from the three *S. pombe* strains were immunoprecipitated using H3K4me3 antibody. The amount of immunoprecipitated DNA was analyzed by qPCR using primer pairs that span promoter of *adh1* sites. The top image shows the adh1 region examined and indicates the position of the primer pairs. Input C_q_ value of each sample was used in normalization. The graph shows the average values of four experiments. Error bars represent the standard deviation. * *P* < 0.05 and ** *P* < 0.01 according to two-sample *t*-test. (**C**) PCA plot of the transcriptomic analyses to visualize the differences between three biological replicates of ADH1 G4 WT, ADH1 G4-TATA MUT, and ADH1 G4 MUT strains. The first principal component (PC 1) accounted for 40.08% and the second principal component (PC 2) accounted for 14.73% of the total variance in the dataset. Volcano plots showing significantly altered transcript levels between three comparison groups: (**D**) ADH1 G4-TATA MUT _vs_ ADH1 G4 WT, (**E**) ADH1 G4 MUT _vs_ ADH1 G4 WT, and (**F**) ADH1 G4-TATA MUT _vs_ ADH1 G4 MUT. Transcripts with significant changes in expression with false discovery rate (FDR) <0.05 and log2 fold change >0.1 are highlighted. Blue and pink dots show down- and up-regulated DEGs, respectively. Eight important genes are marked that are associated to oxidoreduction process. GO pathway analyses bubble plots showing representative top significantly (FDR <0.01) regulated pathways between three comparison groups: (**G**) ADH1 G4-TATA MUT _vs_ ADH1 G4 WT, (**H**) ADH1 G4 MUT _vs_ ADH1 G4 WT, and (**I**) ADH1 G4-TATA MUT _vs_ ADH1 G4 MUT. (**J**) Venn diagram showing common significantly altered genes between the comparison groups associated with oxidoreduction pathway. (**K**) Heatmap profile showing expression (log2(counts per million), scaled by genes) of 8 genes in oxidoreduction pathways that are significantly altered as well as commonly found between three comparison groups.

We also examined the levels of *adh1* antisense RNA transcripts in all three strains, as increased expression of *adh1* antisense RNA under zinc-limited conditions has been reported to repress *adh1^+^*gene expression (91). This regulation is linked to the requirement of zinc for Adh1’s enzymatic activity. However, none of the strains showed elevated *adh1*^+^ antisense RNA transcript levels, suggesting that the reduced *adh1*^+^ transcripts observed in ADH1 G4-TATA MUT and ADH1 G4 MUT strains are unlikely to result from changes in antisense RNA levels (Supplementary Figure 6) (91).

### Adh1 G4 mutations induce minimal alteration in active histone methylation marks

Trimethylation of the fourth lysine residue of the histone H3 protein (H3K4me3) is associated with active transcription (93), and in human embryonic stem cells, promoter G4 structures are often marked with H3K4me3 (33). To determine if the Adh1 G4 mutation impact H3K4me3 distribution across the proximal and distal G4 site in *adh1* promoter, we performed ChIP using H3K4me3-specific antibodies and analyzed the immunoprecipitated DNA by qPCR. Primer pairs were designed to amplify regions spanning the Adh1 G4 site and extending up to the ATG start codon of the *adh1^+^* gene (Supplementary table 2). In general, H3K4me3 marks progressively increased toward the ATG start codon across all strains. However, in the ADH1 G4-TATA MUT strain, which carries mutations in both the Adh1 G4 motif and the conserved TATA sequence, H3K4me3 levels were significantly reduced at the *adh1* promoter compared to ADH1 G4 WT. This reduction aligns with the observed drastic decrease in *adh1^+^* transcription in this strain (Figure 4B). Conversely, the ADH1 G4 MUT strain, which harbors mutations only in the Adh1 G4 motif, showed minor alterations in H3K4me3 levels, primarily near the ATG start codon, relative to ADH1 G4 WT. As expected, no changes in H3K4me3 levels were detected at the *act1* promoter, which served as a non-G4 control site (Supplementary Figure 7). In conclusion, mutations in the Adh1 G4 motif alone caused minimal changes in H3K4me3 distribution, whereas the TATA-box motifs significantly impacted H3K4me3 enrichment at the *adh1* promoter.

### Modification of the dynamics of a single G4 dysregulates cellular transcriptome

Given the association between the Adh1 G4 and transcription, as well as the observed growth impairment in both mutant strains, we performed bulk-RNA sequencing to profile the transcriptomes of all strains. Principal component analysis (PCA) showed distinct clustering of transcriptomes among the different strains, with biological replicates clustering closely together, indicating high reproducibility (Figure 4C). These findings were further supported by Pearson correlation analysis, which confirmed the consistency of transcriptomic data across replicates (Supplementary Figure 8). Volcano plots were generated to identify differentially expressed genes (DEG) between three comparison groups (ADH1 G4-TATA MUT_vs_ADH1 G4 WT; ADH1 G4 MUT_vs_ADH1 G4 WT; and ADH1 G4-TATA MUT_vs_ADH1 G4 MUT) using a fold change >2 and p < 0.05 (Figures 4D - F). A total of 288 and 206 DEGs were identified in the ADH1 G4-TATA MUT_vs_ADH1 G4 WT and ADH1 G4 MUT_vs_ADH1 G4 WT comparisons, respectively. Thus, the mutations in both strains alter the transcriptome of the yeast cells. In the ADH1 G4-TATA MUT_vs_ADH1 G4 MUT comparison, 643 DEGs were identified, with 277 downregulated and 366 upregulated genes (Figures 4D - F), showing also differences between the two mutants. Gene ontology (GO) enrichment analysis highlighted the oxidoreductase pathway as one of the most significantly affected biological processes across all three groups, aligning with the function of the *adh1^+^* gene, which catalyzes the reduction of acetaldehyde to ethanol and the oxidation of NADH to NAD+ (Figures 4G - I). Further analysis of the DEGs in the oxidoreductase pathway identified eight shared DEGs across all three comparison groups (Supplementary Figures 9 - 11, Supplementary Table 3). These eight genes, including *cdc22^+^*, *ilv5^+^*, *nde2^+^*, and *tpx1^+^*, are involved in pathways related to nucleotide metabolism, branched-chain amino acid biosynthesis, the electron transport chain, and the neutralization of reactive oxygen species (ROS) (94–96). These pathways are likely interconnected with the function of Adh1, which plays a key role in maintaining redox balance.

### Modification in G4 dynamics at *adh1^+^* promoter affects the metabolomic profiles in yeast

Since Adh1 plays a central role in redox metabolism and NAD+/NADH balance, mutations in the G4 motif and TATA box of its promoter are likely to affect not only transcriptional regulation (Figure 4), but also downstream metabolic pathways dependent on redox homeostasis. To explore these effects, we extracted metabolites from the three yeast strains and analyzed them by GC-MS. We detected in total 69 metabolites and distinct subsets of differentially abundant metabolites were identified across six biological replicates of ADH1 G4 WT and the two mutant strains (Supplemental data 1). Variability among strains and the consistency between replicates were assessed using Pearson correlation matrix analysis (Supplementary Figure 12). Similar to the transcriptomic analysis, we examined the overlap of differentially abundant metabolites in three comparison groups: ADH1 G4-TATA MUT_vs_ADH1 G4 WT, ADH1 G4 MUT_vs_ADH1 G4 WT, and ADH1 G4-TATA MUT_vs_ADH1 G4 MUT. This analysis showed that 23.4% of the common metabolites were differentially abundant in all three comparison groups (Figure 5A), suggesting intricate relationships among the metabolites affected by the mutations. To further explore the differentially abundant metabolites, we identified those metabolites that were up- or down-regulated in the comparison groups, applying a significance threshold of p < 0.05 and Fold change > 1.2. (Figure 5B - D). Each volcano plot represents the magnitude and significance of differential levels, with metabolites exceeding the thresholds highlighted. This approach provided a clearer understanding of metabolic shifts induced by the mutations and emphasized the overlapping responses among the strains. These included for instance altered levels of niacinamide, citric acid, proline and lysine metabolites. These metabolites are involved in various metabolic and oxidoreductase pathways (97–100). For instance, niacinamide-derived NAD^+^ plays essential roles in redox balance by facilitating electrons transfers between oxidants and reductants (101). Next, we conducted pathway enrichment analyses using these statistically significant metabolites to identify the top 25 significantly impacted pathways resulting from the mutations (Figure 5E - G). In these analyses we observed significant enrichment of amino acid metabolism i.e alanine, aspartate, glutamate metabolism, as well as citric acid (TCA) cycle, and nicotinic acid and niacinamide metabolism in both mutant strains. To gain deeper insights, we conducted pathway impact analysis, which assesses how altered metabolite levels affect pathway functionality and also highlights their critical roles within those pathways (Figures 5H - J). These results further emphasize that the G4 mutation in ADH1 G4 MUT impacts the nicotinic acid and niacinamide pathway, whereas the ADH1 G4-TATA MUT mutation affects multiple pathways, including the TCA cycle, various amino acid metabolism pathways, and the nicotinic acid and niacinamide pathway.

**Figure 5.**
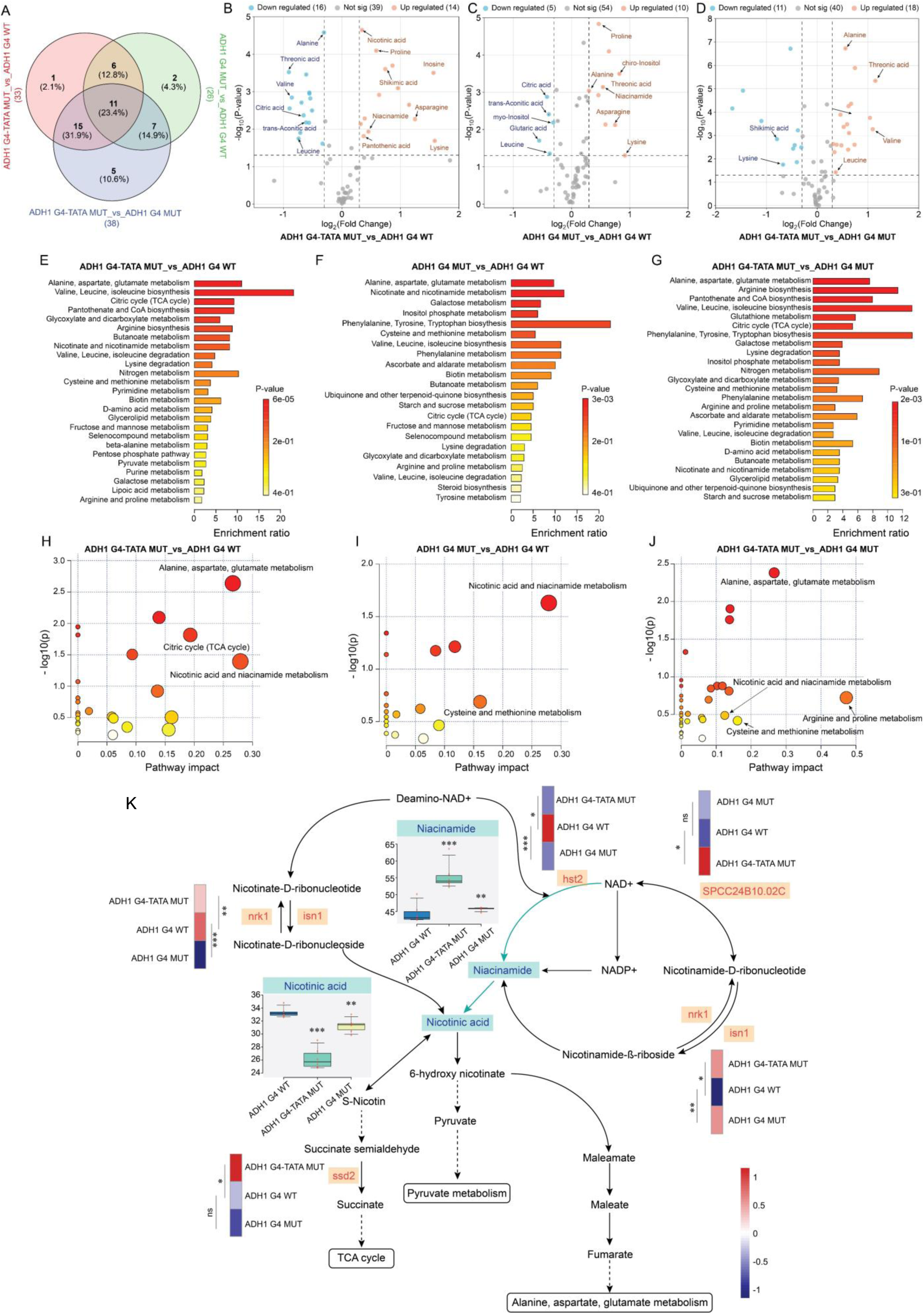
Adh1 G4 motif mutations alter metabolic profiles particularly in nicotinic acid and niacinamide pathways. (**A**) Venn diagram summarizing metabolic changes in ADH1 G4-TATA MUT and ADH1 G4 MUT strains compared to the ADH1 G4 WT strain. Volcano plots displaying metabolome profiles between three comparison groups: (**B**) ADH1 G4-TATA MUT_vs_ ADH1 G4 WT, (**C**) ADH1 G4 MUT _vs_ ADH1 G4 WT, (**D**) ADH1 G4-TATA MUT_vs_ ADH1 G4 MUT with fold change > 1.2 and p-values < 0.05 in metabolite levels induced by Adh1 G4 mutation. Blue and orange dots represent metabolites with significantly decreased and increased levels of the metabolites, respectively, between the comparison groups. Metabolic set enrichment analyses (MSEA) showing top 25 significantly affected Kyoto Encyclopedia of Genes and Genomes (KEGG) metabolic pathways in three comparison groups: (**E**) ADH1 G4-TATA MUT _vs_ ADH1 G4 WT, (**F**) ADH1 G4 MUT _vs_ ADH1 G4 WT, (**G**) ADH1 G4-TATA MUT _vs_ ADH1 G4 MUT using normalized metabolic concentrations from the cells. Color density in the bar plots (white to red) reflects increasing statistical significance. The enrichment ratio is calculated as the ratio of observed hits to expected hits in the pathways. Pathway impact analyses between the comparison groups: (**H**) ADH1 G4-TATA MUT _vs_ ADH1 G4 WT, (**I**) ADH1 G4 MUT _vs_ ADH1 G4 WT, (**J**) ADH1 G4-TATA MUT _vs_ ADH1 G4 MUT, where metabolic pathways are shown as circles based on their −log of p-values from enrichment (y-axis) and the pathway impact values derived from pathway topology analysis (x-axis). Darker circle colors indicate more significant changes of metabolites in their corresponding pathways. The size of the circle corresponds to the pathway impact score and is correlated with the centrality of the involved metabolites. (**K**) Overview of the nicotinic acid and niacinamide metabolism pathway, mostly affected due to Adh1 G4 mutation and its connection to other pathways like TCA cycle, Pyruvate and amino acid metabolism. Box and whisker plots reflecting the altered levels of niacinamide and nicotinic acid between different strains. ** *P* < 0.01 and *** *P* < 0.001 according to paired t-test from six biological replicates. Differential expression levels of DEGs in the oxidoreductase pathways between different strains involved in nicotinic acid and niacinamide metabolism.

Finally, we integrated metabolomic and transcriptomic data to examine the significantly affected nicotinic acid and niacinamide metabolism pathway (Figure 5K). The impact of the G4 mutation was highlighted by key DEGs, such as *nrk1* (Niacinamide Riboside Kinase 1), *isn1* (Inosine-Uridine Nucleoside N-Ribohydrolase 1), and *hst2* (Sirtuin 2 Homolog), which were common to both ADH1 G4-TATA MUT _vs_ADH1 G4 WT and ADH1 G4 MUT_vs_ADH1 G4 WT comparisons. These genes play essential roles in NAD⁺/NADH biosynthesis and metabolism, directly influencing Adh1 activity, which depends on NADH as a cofactor. Furthermore, the TATA box mutation intensified this effect, as evident in specific DEGs such as *SPCC24B10.02C* (NAD/NADH kinase) and *ssd1* (succinate-semialdehyde dehydrogenase), identified exclusively in ADH1 G4-TATA MUT_vs_ADH1 G4 WT. Together, these findings highlight the important role of the *adh1^+^* G4 motif and TATA box in regulating metabolic pathways that contribute to redox homeostasis.

## DISCUSSION

Despite substantial progress in understanding G4s and their broad cellular impacts, the roles of individual G4s are still poorly defined. This gap is particularly evident in unicellular model organisms, where G4 studies remain sparse. Nevertheless, research in systems like fission yeast has historically been instrumental in dissecting cellular regulatory networks (102), suggesting that model organisms offer powerful tools for exploring the specific roles of G4s. In this study, we have established G4 CRISPR-Cas9 approach to introduce systematic site-specific mutations on a G4 motif located upstream of *adh1^+^* gene in fission yeast, *S. pombe*, to understand the individual contribution of a G4 structure in the genome. These mutations allowed us to study the contribution of the G4 structure in regulation of *adh1^+^* gene expression and its downstream effect on yeast metabolism. The *adh1^+^* gene was selected because it is not essential for yeast viability; however, it encodes a key enzyme required for glucose fermentation and contributes to maintaining the NAD+/NADH redox balance (103,104). Genes encoding alcohol dehydrogenases are conserved in eukaryotes and many bacteria, and several paralogs can exist in each organism (105). They have been the focus of attention not only due to their biotechnological relevance in ethanol fermentation, but also medically due to their relevance in alcoholism and alcohol toxicity (106).

Using CRISPR-Cas9, we introduced site-specific point mutations in the G4 motif to precisely alter G4 dynamics while minimizing effects on neighboring chromatin. Unlike previous studies (35–39,107), our mutational approach was more targeted, comparable to CRISPR-based methods used in human cells to explore the role of endogenous *cMYC* G4 folding in regulating *cMYC* oncogene transcription (57). Substituting G with T in G-tetrads, known to disrupt Hoogsteen hydrogen bonding, base-stacking interactions, and K^+^ coordination (108), allowed us to alter G4 dynamics and stability while preserving the structural integrity and broader genomic context of the G4 motif, as confirmed by *in vitro* and *in vivo* analyses.

The dot blot assay showed that the BG4 antibody exhibited higher affinity for ADH1 WT oligonucleotide folded in KCl, while recognition of mutated oligonucleotides was significantly reduced, consistent with previous findings that show that BG4 has high affinity for stable G4 structures (27). CD spectra of mutated oligonucleotides were hypochromic and red-shifted, confirming that G-to-T substitutions altered their topology (83). In agreement, the folded ADH1 WT oligonucleotide migrated faster than its mutated counterparts on a native gel, suggesting that the ADH1 WT oligonucleotide forms compact intramolecular G4 structure, similar to the *S. pombe* telomeric G4 (69). Although the profiles of NMR spectra of the ADH1 WT oligonucleotide differed from those of mutated oligonucleotides, the NMR spectra of the mutated oligonucleotides still displayed imino proton signals between 10-12 ppm, suggesting the possibilities to form G4 involving other Gs in the sequence. The NMR spectra also revealed broadened imino proton signals for ADH1 WT, suggesting polymorphic G4 structures, a feature also seen in human promoter G4s like *cMYC* and *BCL2* (109,110). In contrast, mutated sequences showed discrete imino signals, indicating restricted structural dynamics, similar to those in mutated *cMYC G4* oligonucleotides (Pu22 or Pu24) (111,112), which facilitated their structural characterization. Notably, the ADH1 G4 oligonucleotides (74 nts) are significantly longer than the human G4s studied previously, such as Pu22 (22 nts), highlighting the structural complexity of the ADH1 G4 motif.

We used two independent genome-based assays to investigate G4 structure formation at the *adh1^+^* locus. The qPCR stop assay (71), performed using genomic DNA from *S. pombe* strains with either WT or mutated Adh1 G4 sequences, demonstrated the ability of the ADH1 G4 WT site to form G4 structures. Additionally, ChIP-qPCR with the BG4 antibody confirmed G4 formation, as BG4 specifically recognized and immunoprecipitated Adh1 G4 DNA in the ADH1 G4 WT cells when compared to the corresponding counterparts in the mutated cells. While BG4 has been used in cells from humans and other organisms (21,24,27,28,113), this is its first application in *S. pombe*, establishing it as a valuable tool for G4 analysis in fission yeast. These findings further validate yeast as a robust model for studying the regulatory functions of individual G4s.

We used RT-qPCR and bulk RNA-seq to examine the impact of modifying the upstream G4 structure on *adh1^+^* expression and the transcriptome. Several previous studies have reported G4-mediated upregulation of transcription (30,57,107,114,115). In agreement with this, we observed reduced *adh1^+^* expression following modification of the upstream G4 motif located in the non-coding strand of the *adh1^+^* gene, indicating that the G4 structure positively regulates *adh1^+^* transcriptional activity. In fact, this modification also affected the global transcriptome, primarily altering genes involved in oxidoreduction processes. Supporting this, Pombase, a database for information on *S. pombe* model organism, identifies the molecular function of *adh1^+^* as oxidoreductase activity (https://www.pombase.org/gene/SPCC13B11.01), corroborating its role in oxidoreduction pathway. Dysregulation of oxidoreduction processes in mutated cells could be a downstream event resulting from differential expression of *adh1^+^*. In fact, our metabolomics analysis identified altered abundance of nicotinic acid and niacinamide in both mutants (101), with *nrk1^+^, isn1^+^,* and *hst2^+^* identified as DEGs in both ADH1 G4-TATA MUT_vs_ ADH1 G4 WT and ADH1 G4 MUT_vs_ADH1 G4 WT comparisons. Nrk1 catalyzes the phosphorylation of niacinamide riboside (NR) to form niacinamide mononucleotide (NMN), a precursor for NAD⁺/NADH biosynthesis (116,117). Isn1 supports NAD⁺/NADH synthesis indirectly through nucleotide recycling in the purine and pyrimidine salvage pathways (118). Hst2, a NAD⁺-dependent deacetylase, regulates metabolic processes by deacetylating histone and non-histone proteins, linking NAD⁺ consumption to gene expression and chromatin structure (119–121). As Adh1 depends on NADH as a cofactor for alcohol metabolism, the regulation of NAD⁺/NADH biosynthesis and consumption via Nrk1, Isn1, and Hst2 appears to contribute to maintaining metabolic balance.

We found that CRISPR-Cas9 manipulation introduced two point mutations in the *adh1^+^* TATA box (TATAAATA changed to TAAATATA) in the ADH1 G4-TATA MUT cells. These modifications likely originated from the Cas9-expressing plasmid, where the *cas9* gene is under the control of a mutated *adh1^+^* promoter (adh15) (62) having the same TAAATATA sequence, designed to reduce the expression of the *adh1^+^* promoter. In contrast, the ADH1 G4 MUT strain had precise point mutations exclusively at the G4 site, with the TATA box and neighboring regions unaffected. This demonstrates the accurate targeting of the G4 motif, but also highlights the importance of sequencing surrounding region, especially when targeting gene regulatory regions. Some G4 structures are known to serve as binding hubs for transcription factors, such as SP1 (Specificity Protein 1), which binds G4s in the promoter regions of *cKIT* and *cMYC* oncogenes (122–124). Other transcription factors, including NM23-H2, Pur1, Maz, and PARP-1, also interact with G4 elements in promoters (125–128). The structural dynamics of native G4 motifs often enable the competitive turnover of different transcription factors, with their ability to adopt multiple conformations playing a key role in fine-tuning target recognition and optimal gene expression (129). In this context, a notable example as mentioned above is the *cMYC* G4, where local conformational fluctuations upon protein binding allow the flexible WT G4 structure to transition into more restricted conformations, facilitating distinct biological responses (109,130). We propose that mutations in Adh1 G4 may alter transcription factor binding affinity. While our study, consistent with other recent reports (57,107,115), correlates transcription with G4-forming ability, we cannot rule out the possibility that nucleotide substitutions within G4 motifs might alter B-form DNA, potentially affecting aspects of the transcription machinery. Although G4-binding transcription factors in *S. pombe* remain unidentified, a handful of *S. pombe* G4-binding proteins shown to directly interact with G4 structures have been reported (67,69,76). Future studies will aim to identify transcription factors that interact with G4 structures to clarify their specific contribution in gene regulation.

In summary, our findings indicate that a single G4 structure upstream the *adh1^+^* promoter positively regulates *adh1^+^* transcriptional activity, and its alteration changes the global cellular metabolism. Our findings underscore the power of unicellular model organisms, such as fission yeast, in dissecting the specific roles of G4s within cellular regulatory networks. By employing precise mutational approaches and robust experimental models, we have begun to address the longstanding knowledge gap in understanding the regulatory functions of individual G4s, paving the way for broader insights into their cellular impacts.

## DATA AVAILABILITY

The RNA-seq data are deposited with the GEO accession number GSE285129. To review, go to <u>GEO link</u> and use the following secure token: ylejiyycfdydlen.

## ACKNOWLEDGEMENT

Thanks to Robin Allshire (University of Edinburgh) for providing pLSB-NAT plasmid and constructive feedback for Jesus Torres-Garcia (University of Edinburgh). We also thank Josefin Forslund (Umeå university) for her constructive feedback in the beginning of the project.

We acknowledge the Swedish NMR centre at Umeå University for helping in NMR, and the Protein production Sweden at Umeå University for the expression and purification of BG4. Swedish Metabolomics Centre, Umeå, Sweden (www.swedishmetabolomicscentre.se) is acknowledged for metabolic profiling by GC-MS. We also acknowledge Biochemical Characterization Umeå (BMCU) for access to the CD spectrometer.

## AUTHORS CONTRIBUTIONS

The hypothesis for the study was developed by N.S. and I.O. Experiments were carried out by I.O and P.S. All authors analyzed the data and wrote the manuscript.

