## Supplementary for "CRISPR-Cas9 Targeting of G-Quadruplex DNA in ADH1 promoter Highlights its role in Transcriptome and Metabolome Regulation"

### Supplementary Figures

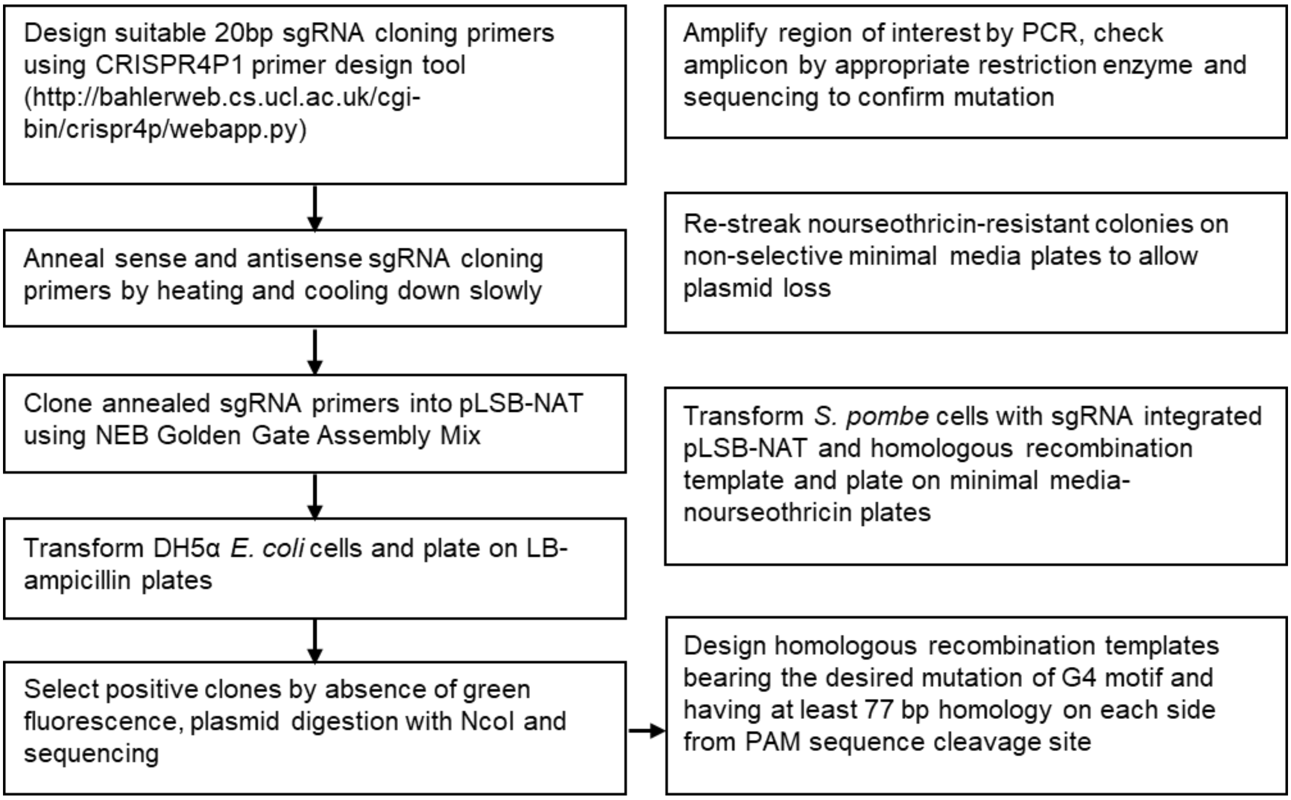

**Supplementary Figure 1.** General flowchart indicating important steps for introducing point mutations in G4 motifs using CRISPR-Cas9 system in *S. pombe*.

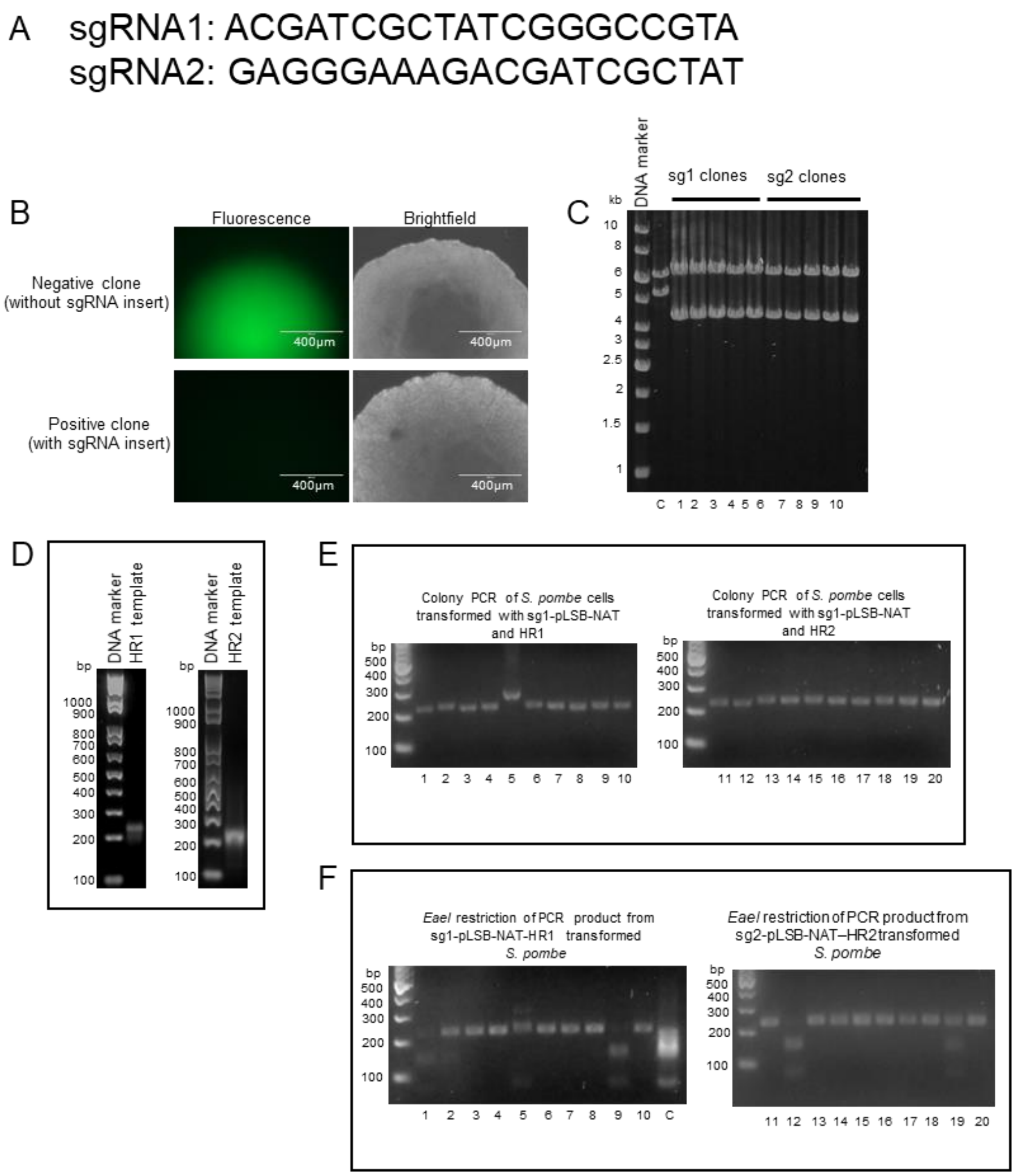

**Supplementary Figure 2.** Generation of *S. pombe* strains with point mutations introduced in the Adh1 G4 motif using G4 CRISPR-Cas9 system. (**A**) Sequences of sgRNA1 and sgRNA2. Sequences (5′-3′) were generated using CRISPR4P1 primer design tool (http://bahlerweb.cs.ucl.ac.uk/cgi-bin/crispr4p/webapp.py) (**B**) Green-white screening of *E. coli* colonies harboring sgRNA inserted into pLSB-NAT plasmid. The presence or absence of GFP was used to screen for negative or positive transformants. Green fluorescent colonies are negative clones without sgRNA insert, while positive clones containing sgRNA insert are without green fluorescence. (**C**) NcoI digestion of pLSB-NAT plasmid isolated from nongreen-fluorescent colonies. The restriction allowed sgRNA-containing plasmids to be distinguished from those containing GFP. Positive clones showed two fragments of correct sizes of ~ 4.3 and 6.3 kb. A control pLSB-NAT plasmid containing only the GFP fragment showed fragments of ~ 5.3 and 6.3 kb. Restricted fragments were analyzed on 0.8% agarose gel. sg1 and sg2 colonies contain sgRNA1 or sgRNA2 integrated in pLSB-NAT, respectively (Lanes 1 – 10). Lane C indicates pLSB-NAT without sgRNA insert. (**D**) Generation of HR templates. Recombinational PCR was used to generate two homologous recombination templates (HR1 and HR2) of 233 bp in size. Products were analyzed on a 2% agarose gel. (**E**) Amplification of 233 bp region containing upstream and downstream regions of Adh1 G4 motif in *S. pombe* cells transformed with sgRNAs-pLSB-NAT and HR templates. (**F**) Screening for *S. pombe* clones harboring Adh1 G4 motif mutations. *Eae*I was used to digest 233 bp PCR products from the G4 site in individual clones. Positive clones showed two fragments of correct sizes of 154 and 79 bp. Fragments were analyzed on a 2% agarose gel. Numbers 1 to 10 denote individual clones of *S. pombe* cells transformed with sgRNA1 integrated in pLSB-NAT and HR1, while numbers 11 to 20 denote individual clones of *S. pombe* transformed with sgRNA2 integrated in pLSB-NAT and HR2. Lane C indicates HR1 DNA, which served as a control.

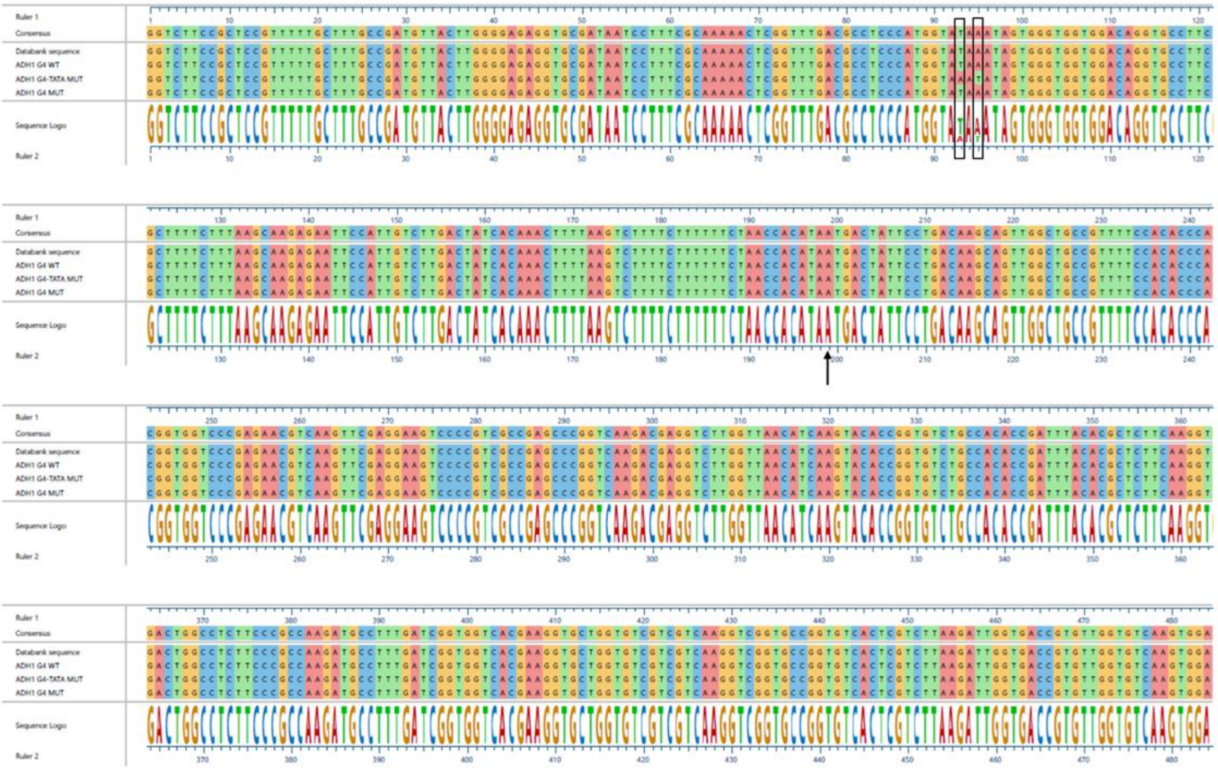

**Supplementary Figure 3.** Sequencing of downstream regions of Adh1 G4. The downstream regions of Adh1 G4 in ADH1 G4 WT, ADH1 G4-TATA MUT and ADH1 G4 MUT *S. pombe* strains were sequenced. The T to A and A to T mutations in ADH1 G4-TATA MUT strain are marked in the boxes. The black arrow denotes the A in Adh1 start codon (ATG).

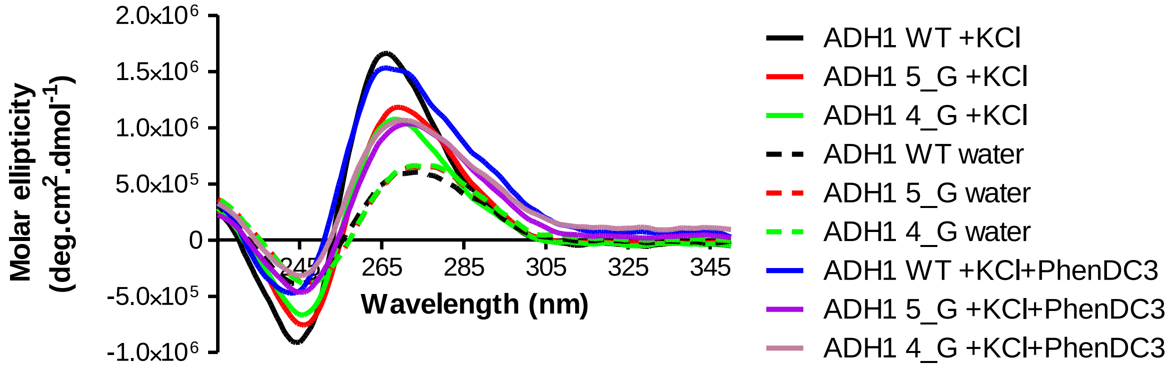

**Supplementary Figure 4.** CD spectra of ADH1 WT, ADH1 5_G MUT and ADH1 4_G MUT oligonucleotides. The oligonucleotides were folded in the presence of KCl and PhenDC$_3$. The data for oligonucleotides folded in only water or KCl are also shown in Figure 3E and are shown here to facilitate comparison with the PhenDC3-treated samples. The CD spectra were recorded between 225 and 350 nm. OriginLab 2020 was used to smoothen the curves.

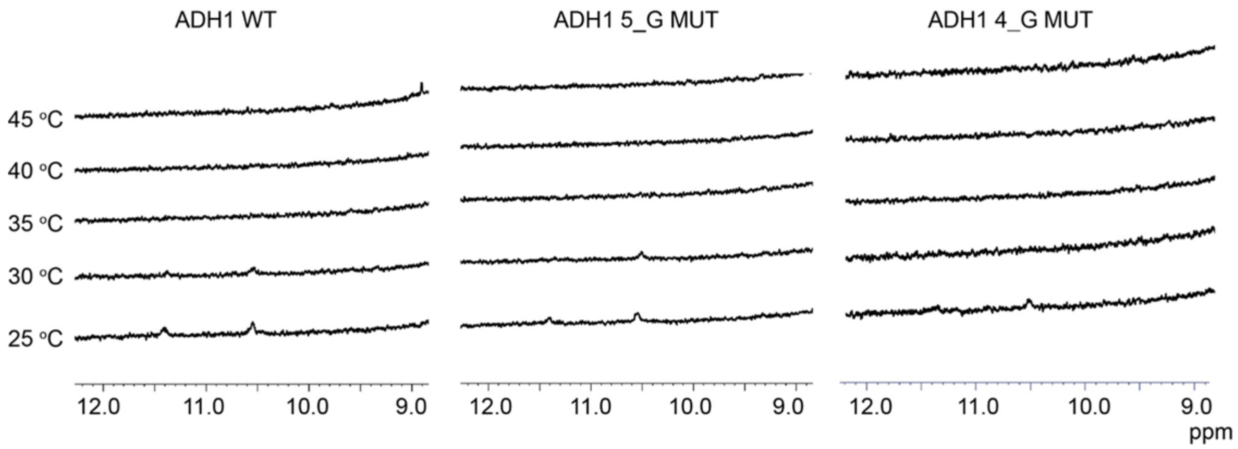

**Supplementary Figure 5.** $^{1}$H NMR spectroscopy of ADH1 WT, ADH1 5_G MUT and ADH1 4_G MUT DNA oligonucleotides in water at different temperatures. The oligonucleotides were folded in water. NMR spectra were recorded at different temperatures on a Bruker 850 MHz Avance III HD spectrometer equipped with a 5 mm TCI cryoprobe and analyzed by Topspin 4.1.3 software.

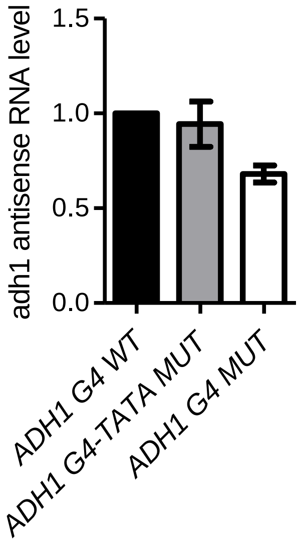

**Supplementary Figure 6.** Realtime qPCR to quantify *adh1* antisense RNA transcript. Total RNA was isolated from ADH1 G4 WT, ADH1 G4-TATA MUT or ADH1 G4 MUT *S. pombe* strains and reverse transcribed to cDNA using a combination of random hexamers and anchored oligo (dT). Primer pairs specific to *adh1* antisense RNA transcript were used.

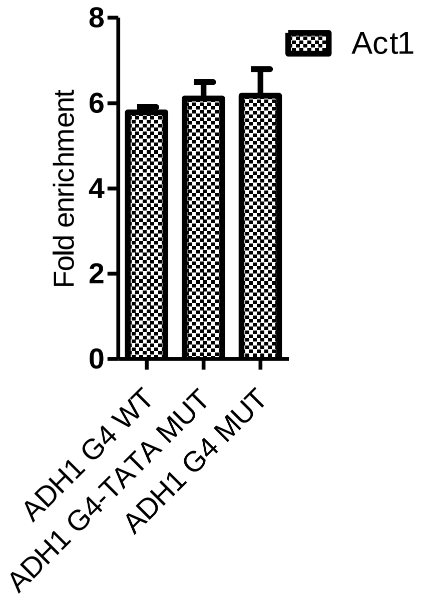

**Supplementary Figure 7.** Assessment of the local chromatin organization on *act1* locus. DNA from chromatin isolated from *S. pombe* strains harboring ADH1 G4 WT or mutated (ADH1 G4-TATA MUT or ADH1 G4 MUT) Adh1 G4 motifs were immunoprecipitated using H3K4me3 antibody. The amount of immunoprecipitated DNA was analyzed by qPCR using primer pairs that span $act1^{+}$ sites. Input $C_q$ value of each sample was used in normalization. The graph shows the average values of four experiments. Error bars represent the standard deviation. * $P < 0.05$ and *** $P < 0.001$ according to two-sample *t*-test.

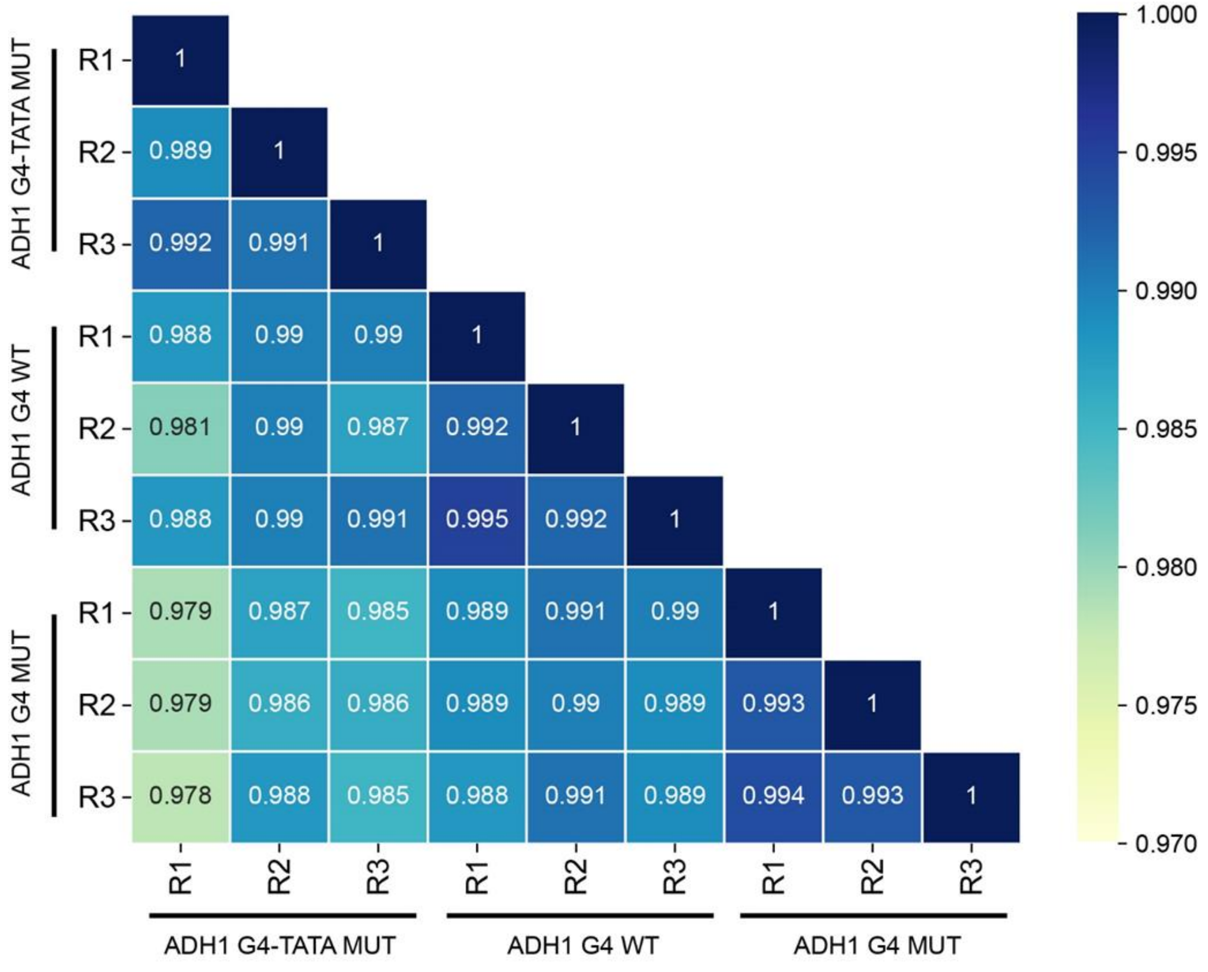

**Supplementary Figure 8.** Pearson's correlation coefficient matrix of bulk RNA-seq between each sample of ADH1 G4 WT and mutants (ADH1 G4-TATA MUT and ADH1 G4 MUT) replicates.

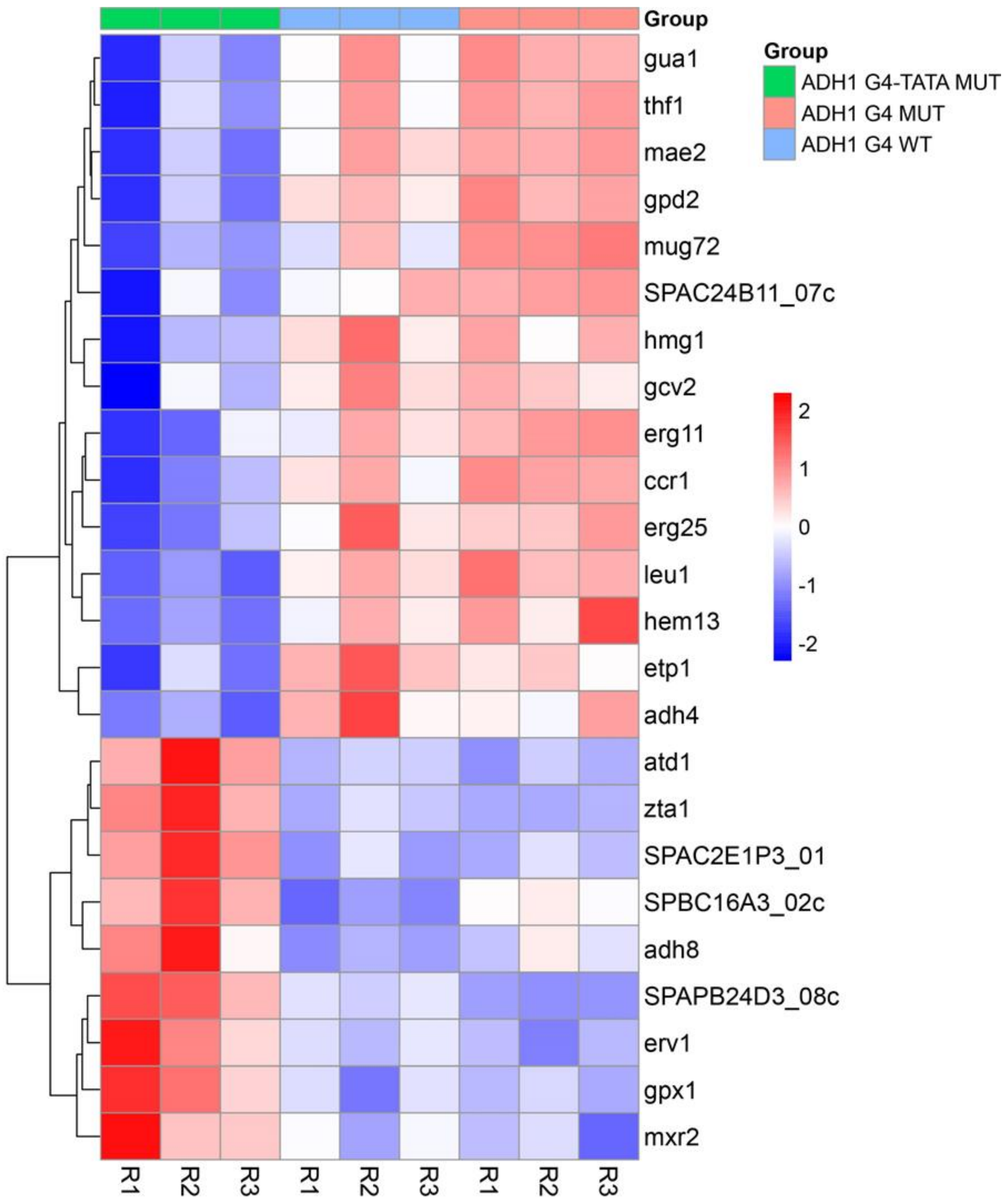

**Supplementary Figure 9.** Heatmap analyses of the 24 significant DEGs in the oxidoreductase pathway that are common between the comparison groups ADH1 G4-TATA MUT _vs_ ADH1 G4 MUT and ADH1 G4-TATA MUT _vs_ ADH1 G4 WT.

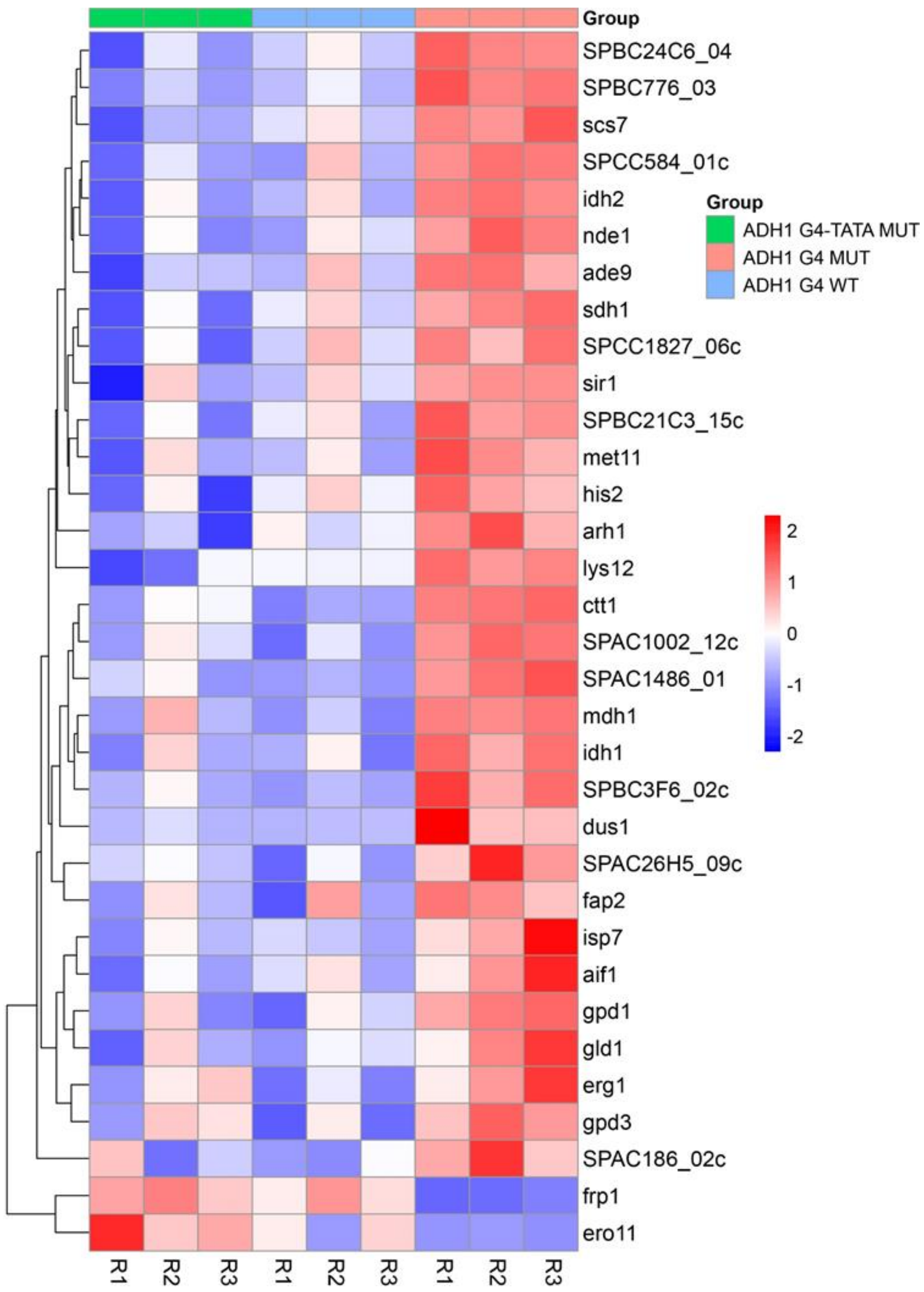

**Supplementary Figure 10.** Heatmap analyses of the 33 significant DEGs in the oxidoreductase pathway that are common between the comparison groups ADH1 G4-TATA MUT _vs_ ADH1 G4 MUT and ADH1 G4 MUT _vs_ ADH1 G4 WT.

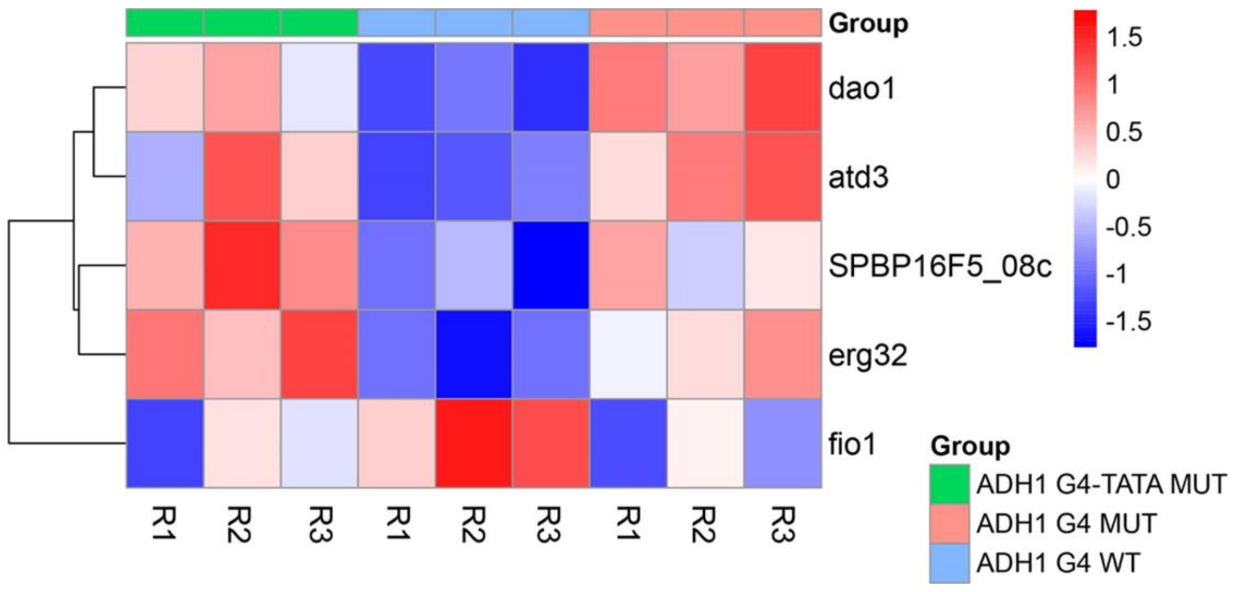

**Supplementary Figure 11.** Heatmap analyses of the five significant DEGs in the oxidoreductase pathway that are common between the comparison groups ADH1 G4 MUT _vs_ ADH1 G4 WT and ADH1 G4-TATA MUT _vs_ ADH1 G4 WT.

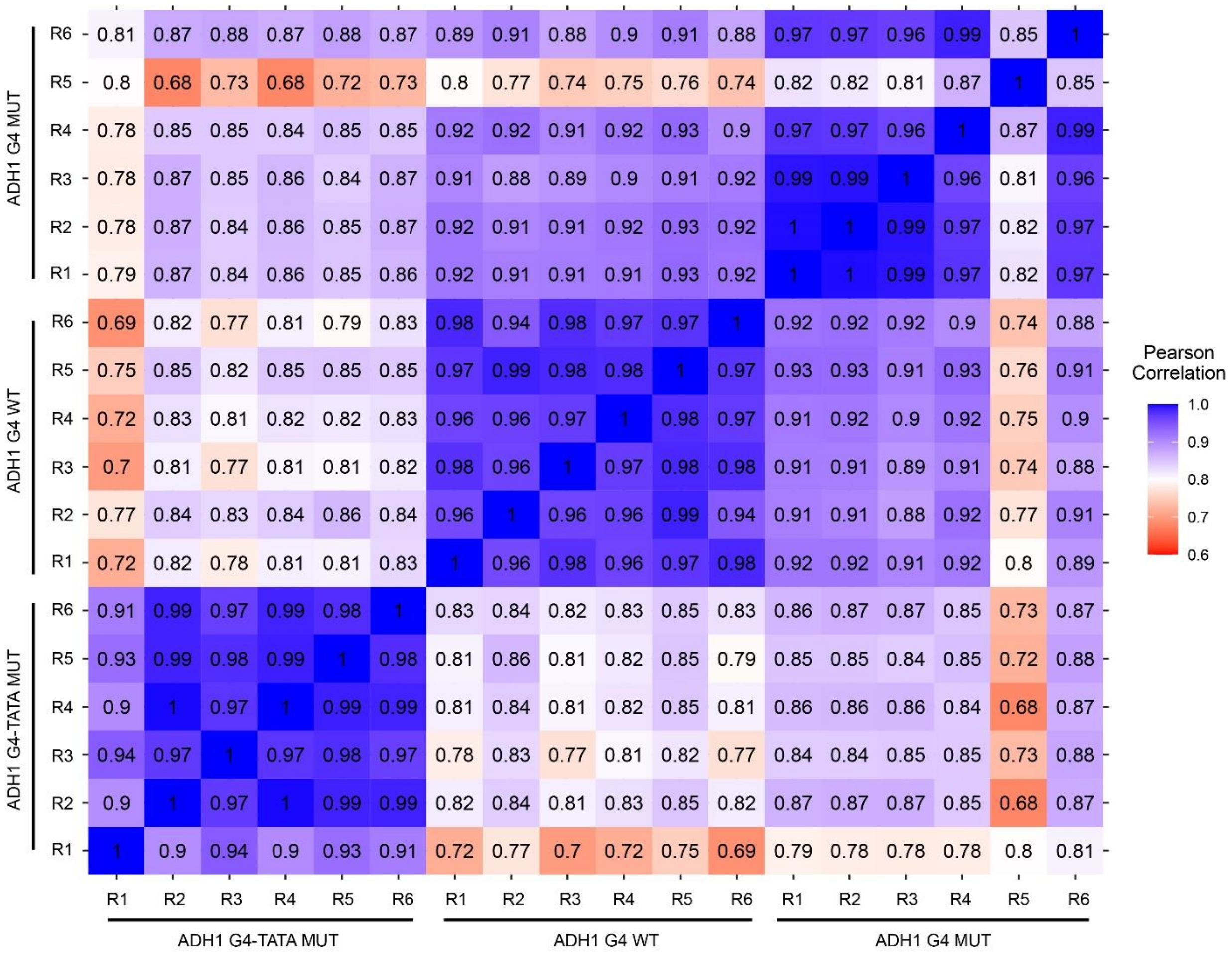

**Supplementary Figure 12**. Pearson's correlation coefficient matrix of metabolomic analyses between each sample of ADH1 G4 WT and mutants (ADH1 G4-TATA MUT and ADH1 G4 MUT) replicates. Six replicates (R1-R6) are shown for each strain.

**Supplementary Tables**

***Strains***

Yeast and bacterial strains used in this study are listed in Supplementary Table 1.

Supplementary Table 1. Yeast and bacteria strains used in this study.

| **Strain** | **Organism** | **Genotype** | **Source** |
|---|---|---|---|
| bNS92 | *E. coli* | DH5α *E. coli* containing pLSB- NAT plasmid inserted with single guide RNA sequence (Sg1) that targets an upstream G4 motif of *adh1* gene. | This study |
| bNS93 | *E. coli* | DH5α *E. coli* containing pLSB- NAT plasmid inserted with single guide RNA sequence (Sg2) that targets an upstream G4 motif of *adh1* gene. | This study |
| YNS119 | *S. pombe* | cdc20+::cdc20-3HA-kanmx6 pfh1::ura4+-nmt-pfh1-GFP ura4-D18 leu1-32 his3-D1 $h^-$ | (1) |
| YIO41 | *S. pombe* | bfr1::hygr pmd::natr cdc20+::cdc20-3HA-kanmx6 pfh1::ura4+-nmt-pfh1-GFP ura4-D18? leu1-32 his3-D1? $adh1_{TATAAATA}$::$adh1_{TAAATATA}$ adh1GGG..GGG..GGG..AGG..GGG::adh1 GTG..GTG..GTG..ACG..GTG | This study |
| YIO43 | *S. pombe* | bfr1::hygr pmd::natr cdc20+::cdc20-3HA-kanmx6 pfh1::ura4+-nmt-pfh1-GFP ura4-D18? leu1-32 his3-D1? | This study |
| YIO45 | *S. pombe* | bfr1::hygr pmd::natr cdc20+::cdc20-3HA-kanmx6 pfh1::ura4+-nmt-pfh1-GFP ura4-D18? leu1-32 | This study |

| | | |
|---|---|---|
| | | his3-D1? adh1GGG..GGG..GGG..GGG::adh1 GTG..GTG..GTG..GTG |

***Oligonucleotides***

Oligonucleotides used in this study are shown in Supplementary Table 2.

Supplementary Table 2. Sequences of oligonucleotides used in this study. The underlined sequences are single guide RNA sequences. The bold letters are the changes in the G-tracks.

| **Oligo name** | **Sequence 5′-3′** | **Description** |
|---|---|---|
| sg1 G435 fw | CtagaGGTCTCgGACTACGATCGCTATCG GGCCGTAGTTTcGAGACCcttCC | Sense strand single guide RNA primer flanked with BsaI sites (marked in blue) for targeting G4 motif upstream of *adh1*$^{+}$ gene |
| sg1 G435 rev | GGaagGGTCTCgAAACTACGGCCCGATA GCGATCGTAGTCcGAGACCtctaG | Anti-sense strand single guide RNA primer flanked with BsaI sites (marked in blue) for targeting G4 motif upstream of *adh1*$^{+}$ gene |
| sg2 G435 fw | CtagaGGTCTCgGACTGAGGGAAAGACG ATCGCTATGTTTcGAGACCcttCC | Sense strand single guide RNA primer flanked with BsaI sites (marked in blue) |

| | | |
|---|---|---|
| | | for targeting G4 motif upstream of *adh1*$^{+}$ gene |
| sg2 G435 rev | GGaagGGTCTCgAAACATAGCGATCGTCTTTCCCTCAGTCcGAGACCtctaG | Anti-sense strand single guide RNA primer flanked with BsaI sites (marked in blue) for targeting G4 motif upstream of *adh1* $^{+}$ gene |
| HR1 G435 mut4 sense | G**T**GGCCGGTAGCGAGTGATAGCGAG**T**GAAAGACGATCGCTATCG**T**GCCGTAACGAGGAAATG**T**GAGGTTGGGGG | Sense strand part of homologous recombination template bearing mutation of the G4 motif of *adh1*$^{+}$ gene |
| HR1 G435 mut4 anti-sense | CCCCCAACCTC**A**CATTTCCTCGTTACGGC**A**CGATAGCGATCGTCTTTC**A**CTCGCTATCACTCGCTACCGGCC**A**C | Anti-sense strand part of homologous recombination template bearing mutation of the G4 motif of *adh1*$^{+}$ gene |
| HR2 G435 mut4 sense | G**T**GGCCGGTAGCGAGTGATAGCGAG**T**GAAAGACGATCGCTATCG**T**GCCGTAAGGAGGAAATG**T**GAGGTTGGGGG | Sense strand part of homologous recombination template bearing mutation of the G4 motif of *adh1*$^{+}$ gene |
| HR2 G435 | CCCCCAACCTC**A**CATTTCCTCCTTACGGC**A**CGATAGCGATCGTCTTTC**A**CTCGCTA | Anti-sense strand part of homologous |

| | | |
|---|---|---|
| mut4 anti-sense | TCACTCGCTACCGGCC**A**C | recombination template bearing mutation of the G4 motif of *adh1*$^{+}$ gene |
| HR G435 upstr. sense | GCGACAGGGGGTCAGCGATAAGCGTCG GAAAAAACTTTATGATATGGTAAATACG TAGGAGGTTACGGTGCAGAGGAGT GGCCGGTAGCGAGTGATA | Sense strand part of homologous recombination template bearing upstream sequences of the G4 motif of *adh1*$^{+}$ gene |
| HR G435 upstr. antisense | TATCACTCGCTACCGGCCACTCCTCTGCA CCGTAACCTCCTACGTATTTACCATATCA TAAAGTTTTTTCCGACGCTTATCGCTGAC CCCCTGTCGC | Anti-sense strand part of homologous recombination template bearing upstream sequences of the G4 motif of *adh1*$^{+}$ gene |
| HR G435 downstr. sense | AGGAAATGTGAGGTTGGGGGAAGGATG CGTTGGAATGCGGAGTAGAGAAATTGAA GAGAGAGAGAGGACACACCTAGAGAGA AAGAAATGGATATAGAGAAA | Sense strand part of homologous recombination template bearing downstream sequences of the G4 motif of *adh1*$^{+}$ gene |
| HR G435 downstr. Anti-sense | TTTCTCTATATCCATTTCTTTCTCTCTAGGT GTGTCCTCTCTCTCTCTTCAATTTCTCTACT CCGCATTCCAACGCATCCTTCCCCCAACCT CACATTTCCT | Anti-sense strand part of homologous recombination template |

| | | |
|---|---|---|
| | | downstream sequences of the G4 motif of *adh1*$^+$ gene |
| HR G435 template fw a | GCGACAGGGGGTCAGCGATA | Forward primer for amplification of 233 bp homologous template |
| HR G435 template rev d | TTTCTCTATATCCATTTCTTTCTCTCT AGG | Reverse primer for amplification of 233 bp homologous template |
| IO20 | GCCAATAGGAGGGCGACAG | Reverse primer for amplification of the G4 motif of *adh1*$^+$ gene |
| IO45 | TCTCTCTAGGTGTGTCCTCT | Forward primer for amplification of the G4 motif of *adh1*$^+$ gene |
| NS213 | CAGTTTAGACGGAAAAGTTTATGC | Forward primer for amplification of *ade6* region |
| NS214 | CACGCTGTTGAATTGAGAAGG | Reverse primer for amplification of *ade6* region |
| IO68 | CCCATGCTACCATCATCCCC | Forward primer for amplification of *adh1* coding region |

| | | |
|---|---|---|
| IO69 | GCCGACCTTGGATTCCTTCA | Reverse primer for amplification of *adh1* coding region |
| IO70 | TGCCAAGCGTGTCATCATCT | Forward primer for amplification of *tdh1* coding region |
| IO71 | GGGGTTGAACTTCTCCTCGT | Reverse primer for amplification of *tdh1* coding region |
| IO72 | TGCTGCTCAATCTTCCTCCC | Forward primer for amplification of *act1* coding region |
| IO73 | TGGAAAAGAGCTTCAGGGGC | Reverse primer for amplification of *act1* coding region |
| IO79 | CGCTTTTGTCCACACCACTC | Forward primer for amplification of *adh1* antisense RNA transcript |
| IO80 | ACCGTCTTTTGACGATTTGGT | Reverse primer for amplification of *adh1* antisense RNA transcript |
| IO94 | CTGCACCGTAACCTCCTACG | Forward primer for amplification of *adh1* promoter region |

| | | |
|---|---|---|
| | | downstream of the Adh1 G4 |
| IO95 | GACAGGGGGTCAGCGATAAG | Reverse primer for amplification of *adh1* promoter region downstream of the Adh1 G4 |
| IO96 | AAATGTGTCCACTTTGGCGG | Forward primer for amplification of *adh1* promoter region downstream of the Adh1 G4 |
| IO97 | CAAAGCAAAAACGGAGCGGA | Reverse primer for amplification of *adh1* promoter region downstream of the Adh1 G4 |
| IO98 | ACAGGTGCCTTCGCTTTTCT | Forward primer for amplification of region proximal to *adh1* start codon |
| IO99 | CAGCCAACTGCTTGTCAGGA | Reverse primer for amplification of region proximal to *adh1* start codon |

| | | |
|---|---|---|
| FO3812 | GAAATCGCAGCGTTGGTTAT | Forward primer for amplification of region proximal to *act1* start codon |
| FO3813 | ACGCTTGCTTTGAGCTTCAT | Reverse primer for amplification of region proximal to *act1* start codon |

Table 3. List of DEGs common to ADH1 G4 WT vs ADH1 G4-TATA MUT, ADH1 G4 WT vs ADH1 G4 MUT and ADH1 G4-TATA MUT vs ADH1 G4 MUT groups in the oxidoreductase pathway.

| **Genes** | **Product** |
|---|---|
| ilv5 | acetohydroxyacid reductoisomerase |
| cdc22 | ribonucleoside reductase large subunit Cdc22 |
| nde2 | external mitochondrial NADH dehydrogenase |
| cao1 | copper amine oxidase Cao1 |
| ofd1 | hypoxic oxygen sensor, prolyl-3,4-dihydroxylase |
| ser3 | D-3 phosphoglycerate dehydrogenase |
| SPBC23G7_10c | Glutathione transporter |
| tpx1 | thioredoxin peroxidase |
